# Modular RNA:DNA nanostructures enable nanopore profiling of ribosomal RNA processing and rRNA variants

**DOI:** 10.1101/2024.08.15.608086

**Authors:** Filip Boskovic, Sarah Sandler, Simon Brauburger, Yuan Shui, Bhavik Kumar, Joana Pereira Dias, Plamena Naydenova, Jinbo Zhu, Stephen Baker, Ulrich F. Keyser

## Abstract

Ribosomal RNAs (rRNAs) serve as species-defining markers and undergo processing steps such as excision of intervening sequences (IVSs). Direct analysis of native rRNAs is hampered by amplification-induced biases and by the high conservation of rRNA sequences, which complicates discrimination of closely related variants. Here, we present modular RNA:DNA nanostructures that enable direct, amplification-free identification of rRNAs and their variants. The approach employs rationally designed RNA:DNA duplexes, named RNA identifiers (IDs), assembled onto native rRNAs *via* short complementary oligonucleotides bearing programmable coding motifs. We demonstrate that native bacterial 16S rRNAs can be directly converted into RNA IDs and detected with solid-state nanopores. Having established direct rRNA readout, we next show that biologically encoded rRNA processing states, including serovar-specific 23S rRNA fragmentation patterns arising from IVS excision, are resolved using RNA IDs. Finally, to extend discrimination beyond processing-level differences, we incorporate catalytically inactive Cas9 ribonucleoprotein complexes to enable single-nucleotide discrimination of rRNA variants. Our modular RNA ID-nanopore system facilitates studying rRNA processing and rRNA diversity.

## INTRODUCTION

Ribosomal RNA (rRNA) is a defining molecular feature of bacteria, functioning both as a core structural component of the ribosome and as an informative marker of species-level identity^1–6^. Bacterial rRNA sequence, processing patterns, and intervening sequence (IVS) presence reflect evolutionary divergence^7–11^. Because rRNA is produced at high copy numbers during active growth^12^, it provides a rich and abundant target for identifying bacterial serovars, rRNA maturation products, and lineage-specific sequence variation. These properties make rRNA an attractive molecule for probing species rRNA diversity and biology directly at the RNA level, rather than through indirect inference from rRNA gene sequences^13,14^.

Although rRNA encodes rich information, experimentally accessing this diversity remains challenging. Many approaches infer rRNA characteristics indirectly from gene sequences or rely on enzymatic amplification steps, which can limit access to native rRNA molecules and their processing intermediates. Enzyme-introduced errors can lead to substantial loss of apparent rRNA diversity, misassigning species as well as introducing quantification errors^3,4,15^. These workflows can mask features such as IVS-containing products, complicate quantitative interpretation of full-length rRNAs, and introduce sequence- or primer-dependent biases^16^. In addition, the large size and complex secondary structure of rRNA pose practical challenges for direct analysis, even with promising long-read sequencing strategies that do not yet provide sufficient fidelity to detect such diversity of native full-length RNA^17–19^. Consequently, complementary approaches that enable direct interrogation of native rRNAs while preserving their processing information would be valuable.

Herein, we introduce modular RNA:DNA nanostructures that enable direct analysis of full-length bacterial rRNAs without reverse transcription and amplification. Short complementary oligonucleotides bearing programmable structural elements are used to reshape native rRNAs into defined RNA identifiers (IDs), which generate characteristic current signatures upon translocation through solid-state nanopores. The ionic current signature of a nanopore translocation reflects the charge, size, and shape of the molecule traversing the pore. The workflow integrates rRNA extraction and RNA ID assembly into a single preparation step while limiting thermal degradation and background RNA. Using this approach, we resolve IVS-containing 23S rRNA processing intermediates from *Salmonella enterica* subsp. *enterica* (*Salmonella)* serovars and detect both rRNA and strain-specific mRNA from *Acinetobacter baumannii*. High specificity is demonstrated by the lack of detectable RNA ID nanopore events when total *A. baumannii* RNA is challenged with hundreds of rRNA oligonucleotides designed for other rRNA species. Incorporation of dCas9–guide RNA complexes further extends the platform to single-nucleotide–resolved discrimination at targeted rRNA variant sites. Together, our approach establishes a generalizable strategy for high-specificity, single-molecule analysis of rRNA IVS processing and sequence heterogeneity.

## RESULTS

### Bacterial full-length rRNA identification with RNA identifiers

We utilized the full-length 16S rRNA transcript and reshaped it using short complementary DNA strands to create an RNA:DNA nanostructure, named an RNA identifier (RNA ID) (Fig. 1a). Selected DNA strands contained overhanging sequences functionalized with biotin, which bind streptavidin and enable sequence-specific structural barcoding (green and red)^20^.

**Fig. 1.**
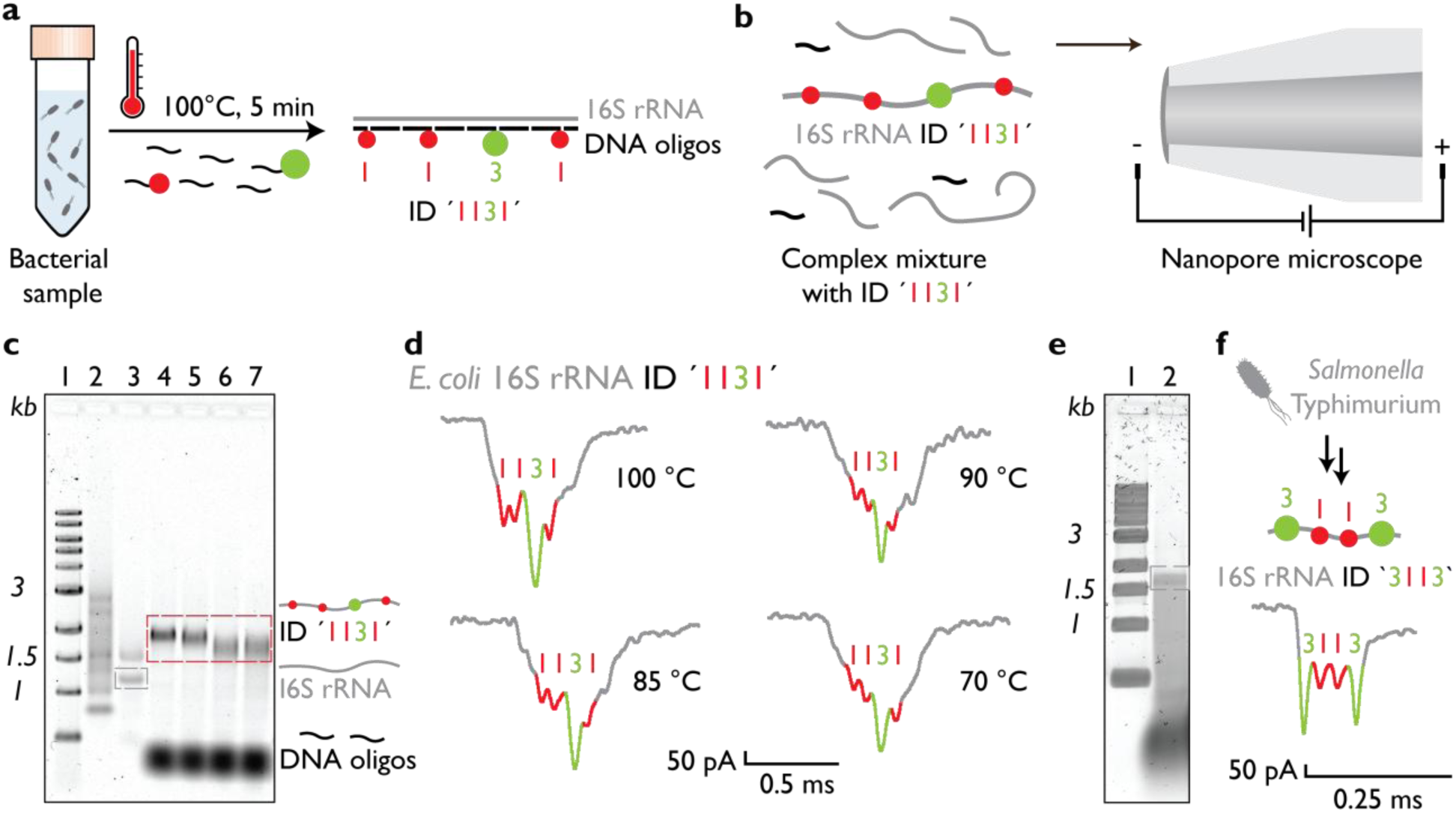
Simultaneous isolation, assembly, and nanopore detection of full-length ribosomal RNA IDs. **(a)** Heating an *E. coli* culture at 100 °C for 5 min in the presence of complementary DNA oligonucleotides enables concurrent rRNA release and assembly of RNA IDs. The 16S rRNA transcript is reshaped into a defined RNA:DNA hybrid duplex through oligonucleotide binding, and placement of docking overhangs at specific positions generates programmable structural codes (e.g., ‘1’ and ‘3’), yielding the *E. coli*–specific 16S rRNA ID ‘1131’. **(b)** Engineered RNA IDs can be detected directly in complex lysates using nanopore microscopy. **(c)** Agarose gel electrophoresis (1% w/v) of *E. coli* total RNA before and after RNA ID assembly shows intact 23S, 16S, and 5S rRNAs prior to treatment (lane 3), whereas samples heated at 70, 85, 90, or 100 °C (lanes 4–7) display selective enrichment of the assembled 16S rRNA ID ‘1131’ accompanied by degradation of unpaired 23S and 5S rRNAs. Lane 1 corresponds to a 1 kbp DNA ladder and lane 2 to an ssRNA ladder. **(d)** Representative nanopore ionic current traces collected over 10 min measurements demonstrate detection of characteristic RNA ID signals for all assembly temperatures. **(e)** Direct assembly of RNA IDs from *E. coli* culture without prior RNA purification yields an approximately 1.6 kb 16S rRNA ID detectable by agarose gel electrophoresis. **(f)** The generality of the approach is demonstrated by assembly and nanopore detection of the 16S rRNA ID ‘3113’ from *Salmonella* Typhimurium culture.

We hypothesized that RNA ID assembly with DNA oligonucleotides could occur during bacterial lysis. To test whether the elevated temperatures required for rapid rRNA release would degrade rRNA during assembly, *E. coli* cells were heated to 75, 85, 90, or 100 °C for 5 min in the presence of DNA oligonucleotides (**Fig. 1a**; sequences listed in Table S1). In this proof-of-concept experiment, *E. coli* 16S rRNA was selected as the target. Our design incorporates both the ‘1’ and ‘3’ structural codes, which denote engineered structural motifs of distinct sizes^20^. Assembly of these elements yields a unique RNA:DNA hybrid duplex, designated RNA ID ‘1131’. The ‘1131’ code also corresponds to the characteristic shape of the nanopore current signal generated by this RNA ID (**Fig. 1b**). This distinct current signature enables detection of 16S rRNA even in complex mixtures containing cellular biomacromolecules and debris such as other RNAs, DNA, lipids, and proteins.

We validated RNA ID assembly using both agarose gel electrophoresis and nanopore measurements by heating *E. coli* DH5α total RNA for 5 min at different temperatures (**Fig. 1c,d**). Prior to treatment, total RNA (lane 3) exhibited three distinct bands corresponding to 23S, 16S, and 5S rRNAs (**Fig. 1c**). After applying the RNA ID assembly protocol to generate 16S rRNA ID ‘1131’ (lanes 4–7), the 23S and 5S rRNA bands were no longer observed, indicating preferential degradation of unpaired rRNAs during the heating step. Successful RNA ID assembly was observed at all four temperatures tested (70, 85, 90, and 100 °C), as evidenced by the remaining band corresponding to the assembled 16S rRNA ID (lanes 4–7). These results indicate selective enrichment of the targeted 16S rRNA ID, highlighted by the red dashed box in **Fig. 1c**.

We next assessed RNA ID assembly at the single-molecule level for all temperature conditions (**Fig. 1d**; additional events shown in **Fig. S1**). In all cases, the characteristic ‘1131’ barcode was detected during a 10 min nanopore measurement. The nanopore results were consistent with the gel electrophoresis observations.

Finally, we applied the heating-based assembly protocol directly to bacterial cultures. *E. coli* samples mixed with DNA oligonucleotides were heated at 100 °C to assemble 16S rRNA ID ‘1131’ (**Fig. 1e**). Agarose gel electrophoresis revealed a band corresponding to the assembled ≈1.6 kb RNA ID (**Fig. 1e**, lane 2). We further demonstrated the generality of this approach using *Salmonella* Typhimurium, targeting its 16S rRNA to generate RNA ID ‘3113’. Single-molecule nanopore events corresponding to this RNA ID were detected within a 10 min measurement (**Fig. 1f;** additional events shown in **Fig. S2**). Together, these results show that RNA extraction, RNA ID assembly, and enrichment can be combined into a single step, enabling efficient identification of full-length bacterial rRNAs at the single-molecule level.

### Discriminating *A. baumannii* variants using rRNA and mRNA RNA IDs

We next applied this approach to *Acinetobacter baumannii*, a bacterium known for extensive genomic and transcriptomic variation, including differences associated with strain-specific genetic determinants^21,22^. One such feature is the presence of the glycosyltransferase gene *gtr100*, which encodes a capsule-associated modification and is present in specific lineages, including KL49 strains^21^. This system therefore provides a useful model to test whether the RNA ID strategy can resolve closely related bacterial strains by simultaneously targeting abundant rRNA and mRNA transcripts.

We designed RNA IDs targeting both the 16S rRNA and the *gtr100* mRNA of a KL49 strain, assembling them into RNA IDs ‘331’ and ‘3113’, respectively (**Fig. 2a**; sequences listed in Tables S2 and S3). Total RNA from two samples, *A. baumannii* lacking *gtr100* (control) and *A. baumannii* KL49 expressing *gtr100*, was analyzed using nanopore measurements (**Fig. 2b**). In both samples, the 16S rRNA RNA ID was readily detected, consistent with the presence of conserved ribosomal transcripts (**Fig. 2c**; additional events shown in **Fig. S3**). In contrast, the RNA ID corresponding to *gtr100* mRNA was detected exclusively in the KL49 sample, reflecting strain-specific transcript expression.

**Fig. 2.**
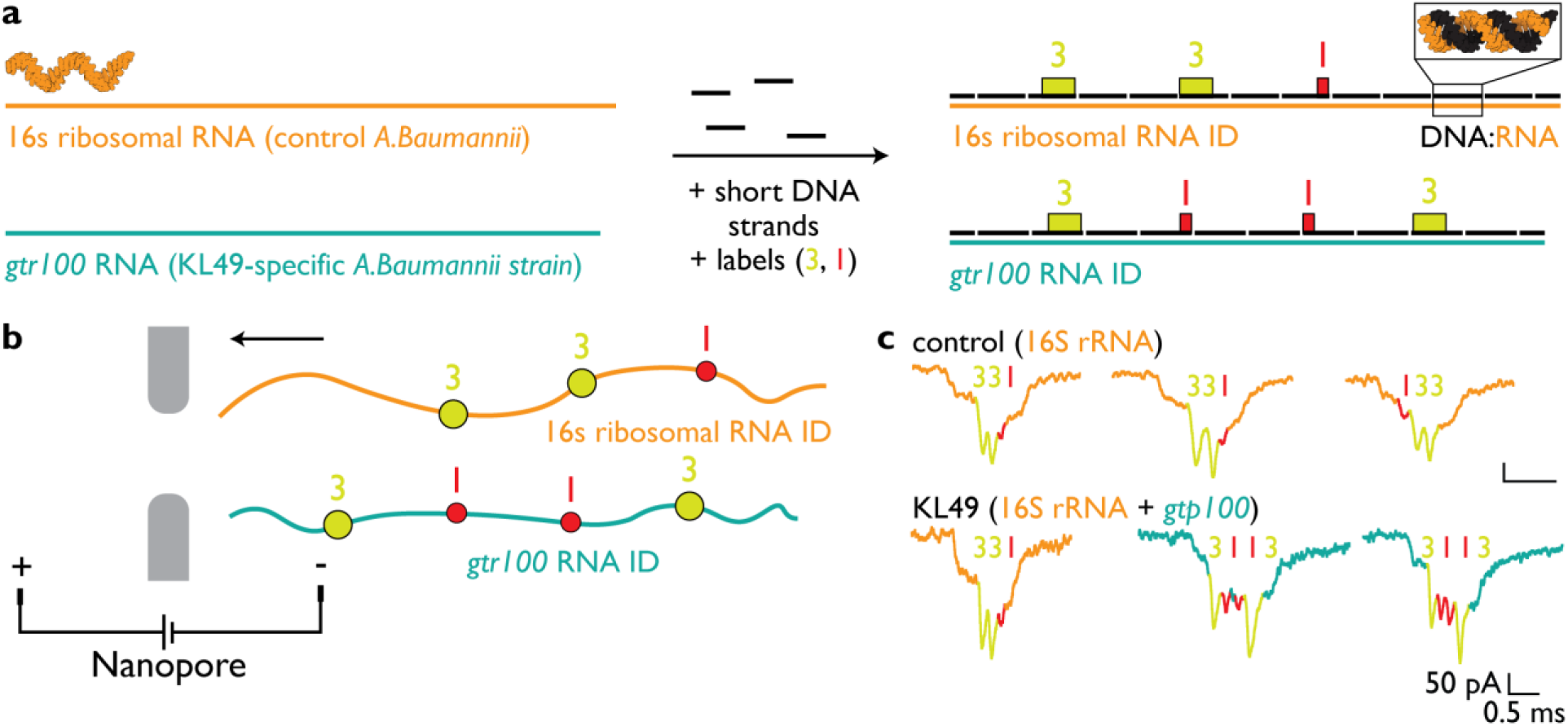
Parallel identification of ribosomal RNA and strain-specific transcript in *Acinetobacter baumannii*. **(a)** RNA IDs were designed to target two RNA species: the 16S rRNA common to *A. baumannii* and the messenger RNA encoded by the strain-associated gene *gtr100*. Assembly of these targets yielded a 16S rRNA ID ‘331’ and a *gtr100* mRNA ID ‘3113’, respectively. **(b)** Schematic representation of nanopore measurements used to analyze RNA IDs assembled from total RNA extracted from two *A. baumannii* strains. **(c)** Representative nanopore events demonstrate detection of the targeted RNA species. The 16S rRNA ID ‘331’ was observed in RNA samples from both strains, whereas the *gtr100* mRNA ID ‘3113’ was detected only in the KL49 strain, consistent with strain-specific transcript expression.

As expected, the *gtr100* mRNA was present at substantially lower abundance than 16S rRNA. Detection of the corresponding gtr100 RNA ID signals occurred at frequencies of approximately 10⁻³–10⁻⁴ per event. Together, these results demonstrate that the RNA ID-nanopore approach can distinguish closely related *A. baumannii* variants by parallel analysis of abundant rRNA and mRNA targets.

### RNA IDs enable multiplexed species identification and co-culture tracking

We evaluated whether RNA IDs could support identification of rRNA transcripts from multiple species or multiplexed tracking within a single experimental framework. Total RNA samples from four phylogenetically distinct sources, human, mouse, *E. coli*, and *Salmonella* Typhimurium, were analyzed to test whether species-specific rRNAs could be selectively converted into RNA IDs (**Fig. S4**). Eukaryotic 18S rRNA and prokaryotic 16S rRNA transcripts were targeted as species-defining RNA molecules (**Fig. S4a**). Four RNA IDs were designed accordingly: ‘1131’ for *E. coli* 16S rRNA, ‘1111’ for human 18S rRNA, ‘3232’ for mouse 18S rRNA, and ‘3113’ for *S.* Typhimurium 16S rRNA (**Fig. S4b**; sequences listed in Tables S1, S4, S5, and S6). Distinct nanopore current signatures corresponding to each RNA ID were observed, demonstrating that multiple rRNA targets can be identified in parallel (additional events shown in **Figs.S1 and S5–S7**).

We further extended this approach to mixed nucleic acid samples containing bacterial and viral components to assess its ability to resolve multiple species simultaneously. RNA IDs were assembled to target *E. coli* 16S rRNA (ID ‘1131’), the single-stranded RNA genome of bacteriophage MS2 (ID ‘111’), and the single-stranded DNA genome of bacteriophage M13 (ID ‘111111’) using a mixture containing *E. coli* total RNA, MS2 RNA, and M13 DNA (**Fig. S8**; sequences listed in Tables S1, S7, S8, and S9). Nanopore measurements revealed distinct current signatures corresponding to each nucleic acid target, indicating successful parallel identification of bacterial rRNA, viral RNA, and viral DNA within a single sample (**Fig. S8**).

To assess the specificity of RNA ID assembly, we challenged the system with a large excess of non-targeting oligonucleotides. A pool of more than 500 oligonucleotides designed to target human 18S rRNA, mouse 18S rRNA, *E. coli* 16S rRNA, and *Salmonella* 16S and 23S rRNAs was mixed with total RNA from *A. baumannii* (**Fig. S9a**). Agarose gel electrophoresis and extended nanopore measurements were then performed to detect any evidence of non-specific RNA ID formation, including the *A. baumannii* 16S rRNA ID ‘331’ (**Figs.S9b and S9c**). Across more than 10 h of measurement time and thousands of nanopore events consistent with RNA or double-stranded DNA molecules, no RNA ID signatures were detected. These results indicate high specificity of RNA ID assembly for the intended target sequences, which we attribute to the combined effects of selective RNA degradation during the heating and assembly step and the distinctive nanopore signals produced by correctly assembled RNA IDs.

### *Salmonella enterica* serovar identification by fragmented 23S rRNA

Having established rRNA-based species identification, we next extended the RNA ID approach to analyze naturally fragmented ribosomal RNA. Several *Salmonella enterica* serovars contain IVS within their 23S rRNA genes with differential combinatorics, which are excised by RNase III during rRNA processing, resulting in defined fragmentation patterns rather than a single full-length transcript^8^ (**Fig. 3a**). Serovars lacking IVSs produce intact 23S rRNA, whereas serovars containing IVS1 or IVS2 generate characteristic sets of rRNA fragments following IVS excision.

**Fig. 3.**
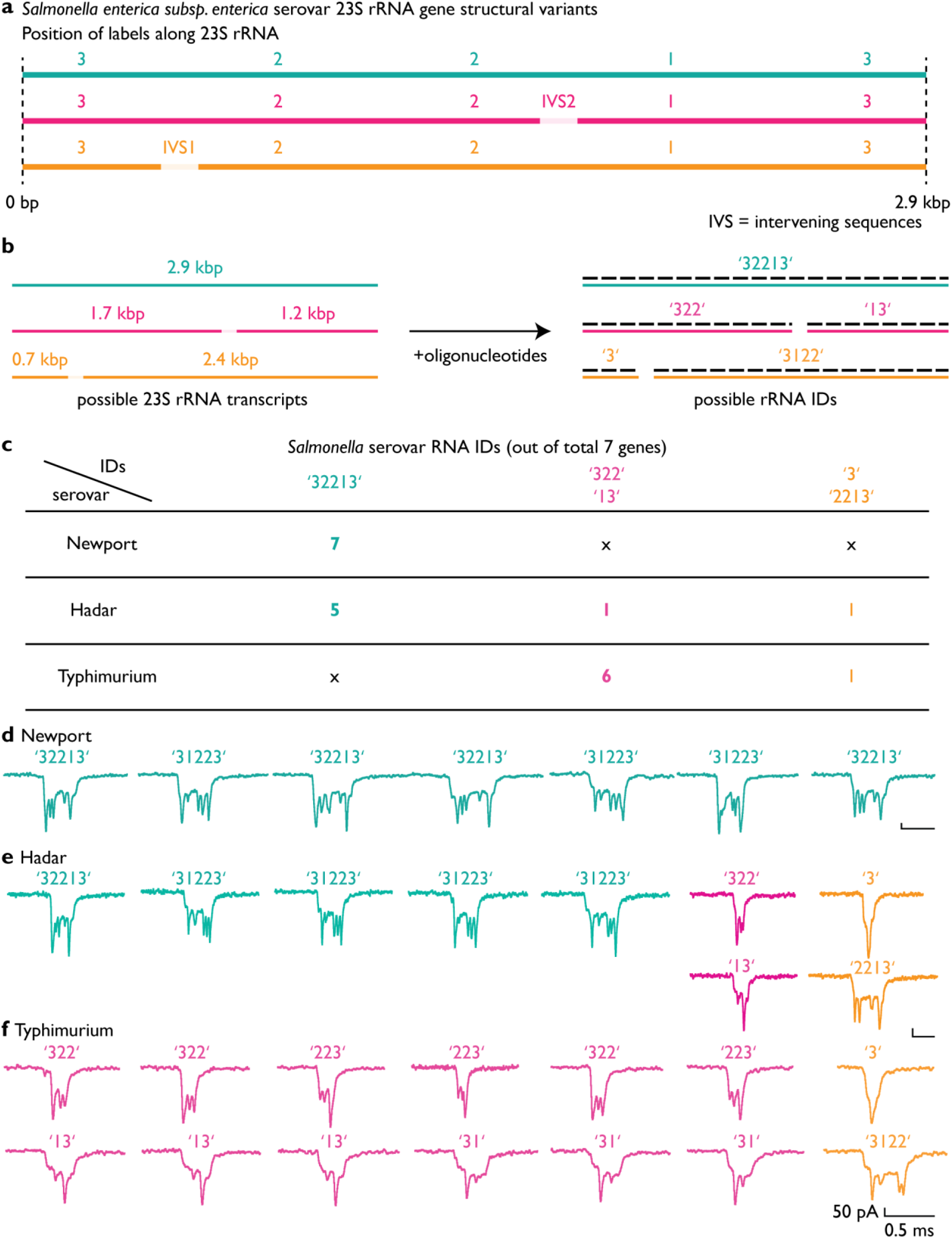
Resolving 23S rRNA processing products from IVS combinatorics using RNA IDs. **(a)** Schematic overview of 23S rRNA processing outcomes arising from different 23S rRNA gene variants. A full-length 2.9 kb 23S rRNA lacking intervening sequences yields the intact RNA ID ‘32213’, whereas variants containing IVS1 or IVS2 undergo RNase III–mediated excision, producing fragment pairs ‘322’ and ‘13’, or ‘3’ and ‘2213’, respectively. **(b)** Assembly of RNA IDs using a single oligonucleotide set generates distinct RNA IDs corresponding to each full-length or processed 23S rRNA. **(c)** Distribution of 23S rRNA gene variants across seven gene copies for the analyzed *Salmonella* serovars, expressed as the number of genes out of seven producing each processing rRNA product. *Salmonella* Newport contains exclusively full-length 23S rRNA genes and yields only RNA ID ‘32213’ (7 of 7), *Salmonella* Hadar contains a mixture of all three variants with predominance of the full-length form (5 of 7), and *Salmonella* Typhimurium contains IVSs in all gene copies and therefore lacks the full-length RNA ID ‘32213’ (0 of 7). **(d)** Representative nanopore events corresponding to 23S rRNA RNA IDs from *Salmonella* Newport. **(e)** Representative nanopore events corresponding to mixed full-length and fragmented 23S rRNA RNA IDs from *Salmonella* Hadar. **(f)** Representative nanopore events corresponding exclusively to fragmented 23S rRNA RNA IDs from *Salmonella* Typhimurium. RNA IDs translocate through nanopores bidirectionally and may therefore be detected in either orientation.

To capture 23S rRNA processing states, we designed an RNA ID architecture that assigns distinct structural codes to full-length and fragmented transcripts (**Fig. 3b**). In this scheme, the full-length 23S rRNA is encoded as ID ‘32213’. Transcripts containing only IVS2 or IVS1 yield fragment IDs ‘322’ and ‘13’, or ‘3’ and ‘2213’, respectively. The expected lengths of all possible ≈2.9 kb 23S rRNA-derived transcripts corresponding to these IDs are summarized in **Fig. 3b**.

We tested this strategy using total RNA from three *Salmonella* serovars, Typhimurium, Newport, and Hadar. Each serovar contains seven 23S rRNA gene copies, which differ in IVS content. In the Typhimurium sample, six genes contain IVS2 and one gene contains IVS1. Using a single oligonucleotide mixture, this composition predicts the presence of fragment IDs ‘322’, ‘13’, ‘3’, and ‘2213’, with no full-length 23S rRNA ID expected (**Fig. 3c**). In contrast, Newport lacks IVSs in all 23S rRNA genes and is therefore expected to predominantly yield the full-length 23S rRNA ID ‘32213’.

Following RNA ID assembly, all three samples were analyzed using nanopore measurements. Representative nanopore events for Typhimurium, Newport, and Hadar are shown in **Fig. 3d, Fig. 3e**, and **Fig. 3f**, respectively. Because RNA IDs can translocate through the nanopore from either end, individual events may appear in either orientation, for example ‘32213’ or ‘31223’ for the full-length 23S rRNA ID.

Consistent with the predicted rRNA processing patterns inferred from their respective genomes, nanopore measurements revealed predominantly full-length 23S rRNA IDs ‘32213’ for Newport (**Fig. 3d**), a mixture of full-length and fragmented IDs ‘32213’, ‘3’, ‘322’, ‘13’, and ‘2213’ for Hadar (**Fig. 3e**), and exclusively fragmented IDs ‘322’, ‘13’, ‘2213’, and ‘3’ for Typhimurium (**Fig. 3f**). These results demonstrate that serovar-specific 23S rRNA processing patterns can be resolved at the single-molecule level using RNA IDs and nanopore readouts. To further extend the resolution of our approach, we next sought to determine whether sequence-level variation within rRNA could also be targeted, motivating the incorporation of dCas9 ribonucleoprotein (RNP) complexes to enable single-nucleotide–specific rRNA discrimination^23^.

### Catalytically inactive Cas9 enables single-nucleotide–resolved rRNA discrimination

To extend RNA ID–based discrimination beyond rRNA processing patterns and into rRNA-level variation, we next incorporated catalytically inactive Cas9 (dCas9) RNP complexes to target single-nucleotide differences within rRNA. dCas9 forms a programmable RNP complex with a guide RNA (gRNA) that binds complementary RNA^24^ in RNA:DNA hybrid context, and under appropriate design constraints can exhibit single-nucleotide specificity. Here, we leverage this programmability to resolve closely related *Salmonella enterica* serotypes based on single-base differences in their rRNA sequences (**Fig. 4**).

**Fig. 4.**
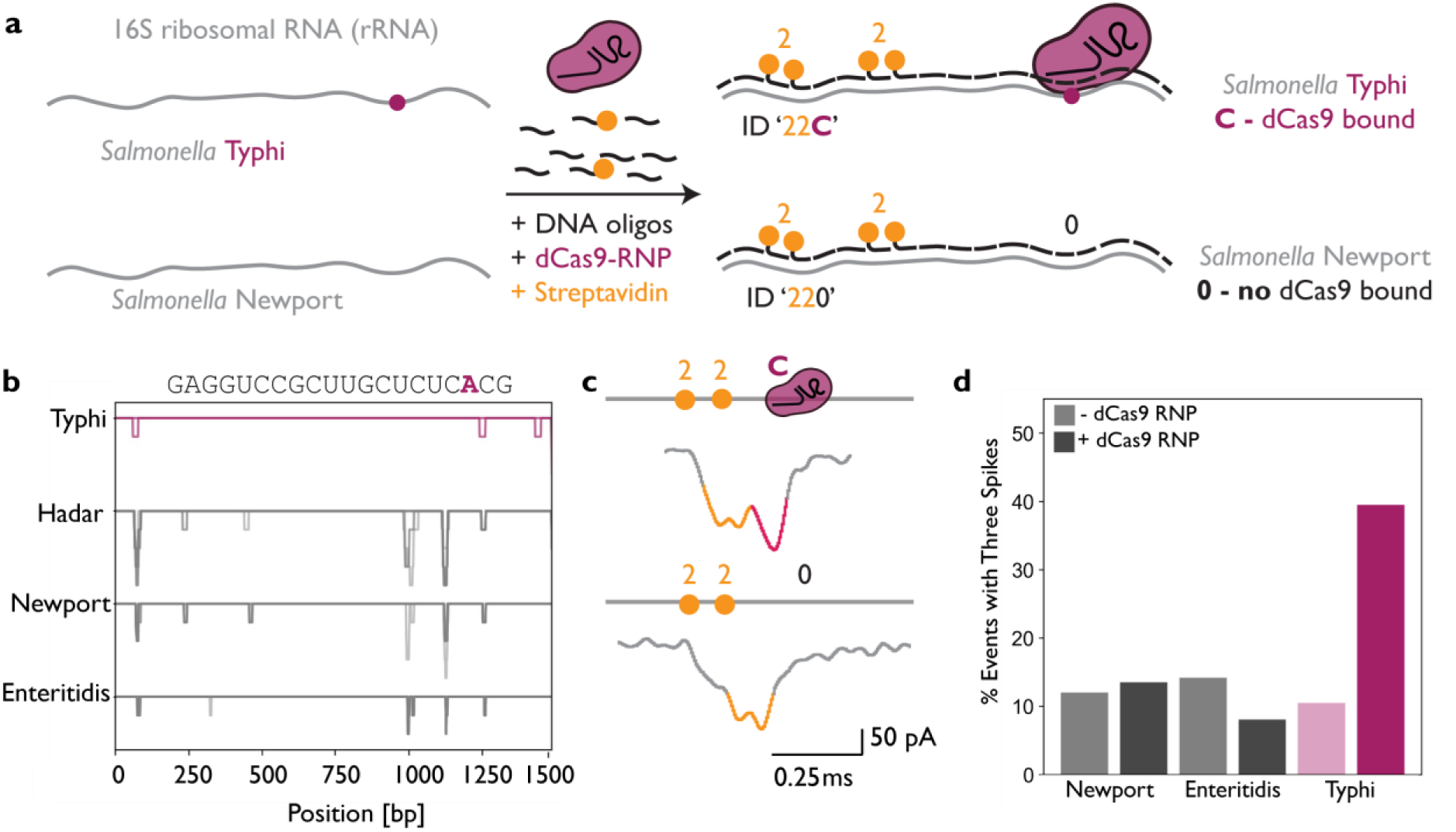
Single-nucleotide discrimination of *Salmonella enterica* rRNA variants using inactive Cas9 (dCas9) binding to RNA IDs. (a) Schematic representation of RNA ID assembly for two closely related 16S rRNA sequences from *Salmonella* serotypes Newport and Typhi that differ by a single nucleotide. The rRNA transcripts serve as scaffolds for complementary DNA oligonucleotides, forming double-stranded RNA:DNA IDs containing defined structural elements corresponding to the bit ‘2’, each associated with two streptavidin molecules bound to biotinylated oligonucleotides. Sequence-specific binding of a dCas9 RNP complex to the Typhi rRNA ID introduces an additional structural element denoted as ‘C’, yielding the barcode ‘22C’, whereas the Newport rRNA ID lacks dCas9 binding and yields the barcode ‘220’. **(b)** Identification of single-nucleotide variant positions by sequence alignment of 16S rRNA from *Salmonella* serotypes Typhi, Hadar, Newport, and Enteritidis, highlighting candidate target sites for dCas9 recognition. **(c)** Representative nanopore events corresponding to RNA IDs ‘22C’ and ‘220’, showing an increased occurrence of three-spike events associated with dCas9-bound RNA IDs in samples containing Typhi. Event counts correspond to measurements performed on samples without Typhi (N = 32) and with Typhi (N = 30). **(d)** Quantification of the fraction of nanopore events containing three peaks before and after addition of dCas9 for Newport, Enteritidis, and Typhi samples. For Newport and Enteritidis, the fraction of three-peak events remains similar with or without dCas9, whereas Typhi shows an increase from approximately 10 percent without dCas9 (N = 45) to approximately 35 percent with dCas9 (N = 33).

In order to use this tool, careful design of the gRNA probes is essential. The binding of dCas9 to RNA:DNA hybrids is constrained by the presence of a protospacer-adjacent motif (PAM) on the DNA strand and by mismatch sensitivity within the gRNA sequence. Previous studies have shown that mismatches proximal to the PAM, particularly within the PAM-proximal “seed” region, most strongly impair Cas9 binding^25^. Variants positioned distally from the PAM often have reduced mismatch sensitivity. Across *Salmonella* serotypes, multiple candidate sites meeting these criteria were identified by 16S rRNA sequence alignment (**Fig. 4b** and **Fig. S10**). Candidate dCas9 RNPs were screened for single-nucleotide specificity using a previously established assay for DNA targeting^23^, and representative measurements are shown in **Fig. S11** (sequences listed in Tables S8, S10, and S11). From these candidates, the gRNA targeting a single-nucleotide variant present at the highest frequency among 16S rRNA gene copies was selected for further analysis.

The selected gRNA which was used for an initial demonstration, allows us to differentiate *Salmonella* Typhi and Newport, which differ by a single nucleotide at position 1258 of the 16S rRNA (**Fig. 4a,b**). A dCas9 RNP was assembled with a gRNA complementary to the Typhi sequence at this position. Binding of the dCas9 RNP to the 16S rRNA RNA ID introduces an additional structural feature, which is encoded as the barcode ‘22C’, where ‘C’ denotes the presence of bound dCas9 and each numeric element corresponds to streptavidin-associated structural features as described above. A unique dCas9 nanopore signature has been demonstrated for dsDNA^23,26,27^, and similarly when dCas9 is bound on RNA:DNA, ‘C’ is detectable. In contrast to Typhi, the Newport 16S rRNA contains a single-base substitution at the target site, preventing dCas9 binding and yielding the barcode ‘220’. Representative nanopore events corresponding to these two RNA ID configurations are shown in **Fig. 4c**, demonstrating discrimination between the two rRNAs based on a single-nucleotide difference.

Using this optimized dCas9 RNP, we compared 16S rRNA RNA IDs assembled from *Salmonella* Typhi, Enteritidis, and Newport (assembly sequences for RNA ID ‘022’ listed in Table S12). Nanopore measurements revealed a higher relative frequency of RNA ID events containing the ‘22C’ barcode in the Typhi sample relative to the other serotypes, which predominantly yielded ‘220’ events lacking dCas9 binding (**Fig. 4d**). Independent repeats of RNA ID assembly and nanopore measurement for Typhi produced consistent results (**Fig. S12**).

Finally, we tested whether dCas9-mediated discrimination could be maintained in mixed-serotype samples. Two mixtures were analyzed, one containing Enteritidis and Newport only, and a second containing Enteritidis, Newport, and Typhi (**Fig. S13a**). Nanopore events were classified based on the presence (’22C’) or absence (’220’) of dCas9 binding, with representative events shown in **Fig. S13b**. Consistent with expectations, ‘22C’ events were observed only in mixtures containing Typhi, demonstrating that dCas9-enabled RNA IDs can resolve single-nucleotide rRNA variants even in the presence of closely related sequences.

## Discussion

By converting native rRNAs into programmable RNA IDs readable by solid-state nanopores, we demonstrate that abundant rRNA products are reshaped into distinct, information-rich signals without reverse transcription and amplification. By reshaping full-length rRNAs into RNA IDs with defined structural signatures, we enabled single-molecule rRNA detection, 23S rRNA processing state specific to *Salmonella* serovars, and 16S rRNA sequence variation specific to serovars with dCas9 RNP binding.

Solid-state nanopores have been widely applied to the analysis of DNA^28–32^, viruses^33–35^, proteins^36–39^, and RNA^20,36,40–43^, including studies of complex biological assemblies^44,45^. Previous nanopore-based RNA measurements have typically relied on purified targets or have been limited in specificity when applied to heterogeneous samples^41,43,44^. The RNA ID builds on these strengths by introducing programmable structural encoding^20^, which enables selective barcoding of RNAs of interest and discrimination from background nucleic acids within complex mixtures. Direct nanopore analysis of native RNA is especially valuable for studying RNA variants and isoforms that differ in length or structure, features that are difficult to preserve through enzymatic workflows^20,46–48^. At the same time, RNA:DNA duplexes translocate faster than DNA:DNA duplexes^49^, highlighting the importance of continued development of velocity-control strategies to further expand nanopore spatial resolution^50–53^.

Ribosomal RNA processing is inherently complex and variable across species, in part because rRNAs are encoded by multiple gene copies that can undergo distinct processing pathways^1,7,10,13^. As a result, a single cell can contain a diverse population of rRNA products arising from differential cleavage, excision of IVS, and regulated maturation steps. These processes are increasingly recognized as responsive to genetic context and cellular conditions^13,54–57^, underscoring the importance of accessing full-length, native rRNA populations. The RNA ID approach provides a complementary route to studying 16S and 23S rRNA transcripts by facilitating direct readout of rRNA products without enzymatic modifications of the RNA itself. This is particularly relevant for transcripts that are highly structured or modified, which may be inefficiently reverse-transcribed or underrepresented in sequencing-based workflows unless specifically accommodated by experimental design^58,59^.

Sequence-level discrimination within rRNA remains challenging due to the high conservation of ribosomal sequences^3,4,15,18,60^. Incorporation of dCas9 RNP complexes adds an additional, programmable recognition layer to the RNA ID framework, enabling single-nucleotide–resolved discrimination within otherwise similar rRNAs. dCas9-based targeting has previously been employed for specific RNA detection^24,61,62^, and its application herein to RNA:DNA hybrid nanostructures demonstrates a promising route for introducing sequence selectivity without altering the underlying RNA ID architecture. Such an approach is particularly relevant for distinguishing closely related bacterial serovars, where single-nucleotide differences within rRNA can carry important biological meaning^63^. However, the applicability of a dCas9-based approach is constrained by the requirement that the distinguishing nucleotide lie sufficiently close to a protospacer-adjacent motif (PAM) to enable selective dCas9 binding. Future extensions of the RNA ID framework could incorporate alternative programmable RNA-or DNA-binding proteins that do not rely on PAM recognition, potentially expanding the range of accessible sequence variants.

While our study focuses on capabilities for rRNA detection, the abundance of rRNA and the ability to resolve species-, serovar-, and variant-level differences suggest potential relevance for future diagnostic applications. Translation to clinical settings will require careful consideration of sample handling, RNA stability, and quantitative interpretation in complex biological environments. Our work provides a route for converting native rRNA molecules into programmable nanostructures, paving the way for single-molecule studies of rRNA processing and rRNA heterogeneity.

## Materials and Methods

### Salmonella enterica and Acinetobacter baumannii cultures and total RNA isolation

Bacteria were cultured overnight in LB and 500 µL of overnight culture was treated with two volumes of RNAprotect reagent (Qiagen). Cultures were digested with 15 mg/mL lysozyme (Sigma) for 20 min and RNA extraction was performed using the RNeasy Mini Kit (Qiagen) following the manufacturer’s guidelines. Total RNA was quantified, and quality assessed using Nanodrop (ThermoFisher). Total RNAs from other samples were obtained from ThermoFisher.

### RNA identifier assembly

RNA IDs were assembled using 5-10 µL of total RNA (ng/µL), 2.4 µL of oligopool (Integrated DNA Technologies; concentration of each oligo is 1 µM), 4 µL of 1 M LiCl, 4 µL of 10 × TE (100 mM Tris-HCl pH 8.0, 10 mM EDTA), and 1 µL of 25 µM 3’-biotinylated imaging strand (IDT; 5’-ACCACTAATGAGTGATATCC-TEG-biotin-3’), nuclease-free water was added to a final volume of 40 µL^26^. Samples were heated to 70 °C for 30 s, slowly cooled to 20 °C over 45 min, and stored at 4 °C or filtered immediately (ProFlex PCR System (Applied Biosystems)). The 40 µL mixture was mixed twice with 460 µL of a washing buffer (10 mM Tris-HCl pH 8.0, 0.5 mM MgCl_2_) and centrifuged in 0.5 mL Amicon ultra centrifugal filter 100 kDa (Sigma-Millipore, catalogue number UFC510008) at 9,400 × g for 10 min. The filtered RNA IDs were retrieved by inverting the filter to a new 2 mL tube and centrifuged at 1,000 × g for 2 min and stored at 4 °C prior the use. All buffers were filtered using Millipore 0.22 µm syringe filters.

### Short heating protocol for RNA ID assembly

RNA ID was assembled using the identical procedure except the heating step as mentioned for RNA ID. The heating protocol was changed to 70, 80, 90, or 100 °C for 5 min, placed on ice for 5 min before ultrafiltration with Amicon 0.5 mL filter.

### DNA carrier assembly

The DNA constructs contained different “barcodes” plus a customizable overhang sequence that is a DNA version of the target RNA sequence. The DNA construct was synthesized from pairing a linearized 7.2 kbp single-stranded (ss) M13mp18 DNA (Guild Biosciences) and with 190 complementary oligonucleotides via Watson-Crick base pairing to produce full dsDNA for 1 h in a thermocycler. All oligonucleotides were synthesized by IDT and dissolved in IDTE (10 mM Tris-HCl, 0.1 mM EDTA, pH 8.0). The sequences are listed in Table S1^52^. The sample is then filtered using a 100 kDa Amicon filter and measured with a ThermoFisher Scientific Nanodrop™ 2000 Spectrophotometer for concentration information. All samples were stored in a storage buffer of 10 mM Tris-HCl 0.5 mM MgCl_2_ after ultrafiltration. The design contains five groups of equally spaced simple dumbbell hairpin motifs within the 190 oligos to create the spikes which act as a barcode on the DNA nanostructure. Each group consists of 11 DNA dumbbells to create a single spike^52^. The sequences modified to contain DNA dumbbells are listed in Table S2. The overhang used for the target was created by replacing oligos No. 142 and 143 from Table S1 with 50 bp target sequences as shown in Table S3.

### CRISPR-dCas9 and streptavidin binding assay

Catalytically deactivated Cas9 D10A/H10A (dCas9) from *Streptococcus pyogenes* binds with a tracrRNA and a sequence-specific RNA (crRNA), both synthesized by Integrated DNA Technologies (IDT) and dissolved in IDTE (10 mM Tris–HCl, 0.1 mM EDTA, pH 8.0). The target sequences for the crRNA for the probes were designed using online software (http://chopchop.cbu.uib.no/). To assemble the dCas9 RNPs, first the tracrRNA (200 nM) was incubated at 90 °C for 2 minutes in a 1× low salt buffer (25 mM HEPES-NaOH (pH 8.0), 150 mM NaCl, 1 mM MgCl_2_) and then quickly cooled by placing the tube on ice. After 3 minutes, the dCas9 (100 nM) was added and allowed to sit at 25 °C with tracrRNA for at least 5 minutes. After 5 minutes, the crRNA (250 nM) was added and allowed to sit for at least 10 minutes at 25 °C. Finally, the assembled dCas9 probes were then incubated with the DNA nanostructures or RNA-IDs for at least 20 minutes at 25 °C, with the dCas9 probes added in excess of typically 15 dCas9 probes per DNA binding site. For the RNA IDs, after dCas9 binding, streptavidin was added in 10 × excess per binding site directly before measurement and allowed to sit at 25 °C for 10 minutes. The samples are then diluted to 0.1 nM DNA nanostructure or RNA ID into a 2 M LiCl, 1 × TE buffer solution immediately before the beginning of the measurement in the nanopore system.

### Agarose gel electrophoresis

Our RNA ID constructs were run on a 0.8-1% (w/v) agarose gel (agarose for molecular biology, Sigma, catalogue number A9539), 1 × TBE (0.089 M Tris-Borate, 0.002 M EDTA, pH 8.3; Sigma, catalogue number SRE0062) in an ice bath, for 1.5 h at 70 V. The gel was poststained in 3 × GelRed solution (Biotium, catalogue number 41001) for 10 min on a horizontal shaker. The stained gel was imaged with a GelDocIt™(UVP).

Gel images were analyzed using ImageJ (Fiji)^55^ by inverting grayscale and homogeneous background subtraction with 100-pixel rolling ball.

### Nanopore measurements

Conical nanopores with diameters of 10-15 nm are made of commercially available quartz capillaries (0.2 mm inner diameter/0.5 mm outer diameter Sutter Instruments, CA). They are created using a laser-assisted pipette puller (P-2000, Sutter Instrument, CA). Sixteen nanopores are then placed in a custom-templated polydimethylsiloxane (PDMS) chip containing a communal cis reservoir and individual trans reservoirs. In order to generate the current, silver/silver chloride (Ag/AgCl) electrodes are connected to the cis and trans reservoirs in the PDMS chip. As only one nanopore can be measured at a time due to the electronics, the trans reservoir contains the electrode with a 500 mV bias voltage, and the central cis reservoir is connected to a ground.

Using an Axopatch 200B patch-clamp amplifier (Molecular Devices, CA), current signals are collected. The setup is operated in whole-cell mode with the internal filter set to 100 kHz. An 8-pole analogue low-pass Bessel filter (900CT, Frequency Devices, IL) with a cutoff frequency of 50 kHz is used to reduce noise. The applied voltage is controlled through an I/O analog-to-digital converter (DAQ-cards, PCIe-6251, National Instruments, TX). A LabView program records the current signal at a bandwidth of 1 MHz.

### Nanopore data analysis

For data using DNA carrier and dCas9 the following procedure was followed. First, a translocation finder python script (https://gitlab.com/keyserlab/nanopyre) is used that identifies the events from the raw traces using user-defined thresholds (minimum 0.1 ms duration, minimum 0.1 nA current drop) and stores them in an hdf5 file. Next, the hdf5 file is loaded into the GUI categorizer python script, found here: https://gitlab.com/keyserlab/nanopycker. Events are then sorted based on the number of peaks or the barcodes and presence or absence of proteins bound.

The binding of dCas9 to RNA:DNA *Salmonella* samples:

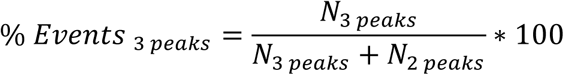

Where three peaks correspond to ‘22C’ ID and 2 peaks correspond to ‘22’ ID. Events containing one peak or no peaks were discarded.

The specificity as seen in Fig. S11. was calculated using the following formula:

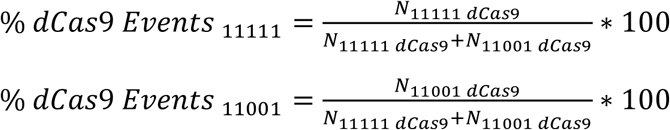

## Acknowledgements

F.B. acknowledges research funding from the George and Lilian Schiff Foundation Studentship, the Winton Programme for the Physics of Sustainability PhD Scholarship and St John’s College Benefactors’ Scholarship. S.E.S. acknowledges funding from Oxford Nanopore Technologies, Engineering and Physical Sciences Research Council (EPSRC) and Cambridge Trust. U.F.K. acknowledges funding from a European Research Council (ERC) consolidator grant (DesignerPores no. 647144) and an ERC-2019-PoC grant (PoreDetect no. 899538).

