## Supplementary Materials for "Modular RNA:DNA nanostructures enable nanopore profiling of ribosomal RNA processing and rRNA variants"

**This document includes**

Figs. S1 to S13
Tables S1 to S15

**Fig.S1**

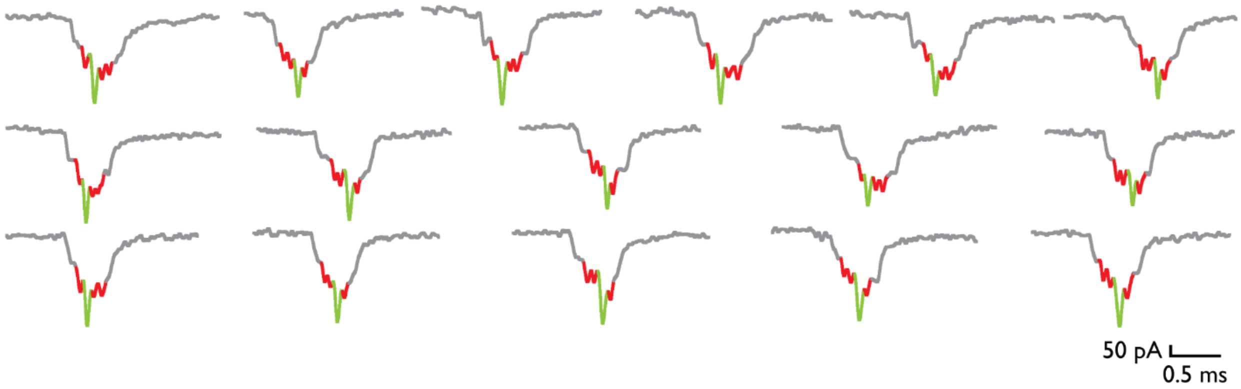


**Fig.S1** Typical nanopore events for *E.coli* J01859.1 16S rRNA ID `1131`.

**Fig.S2**

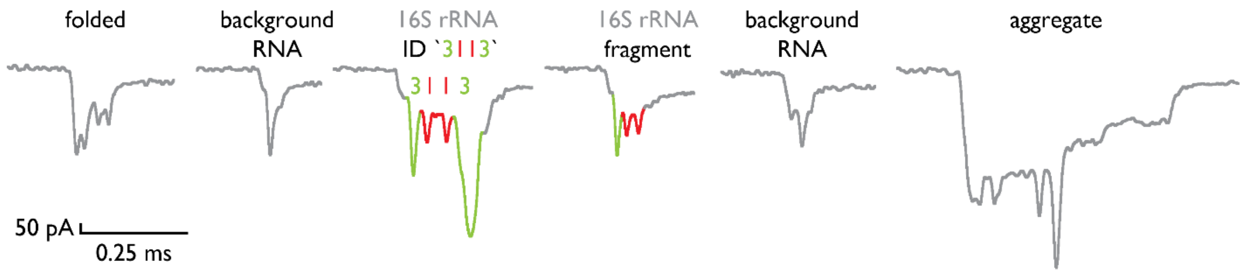


**Fig.S2** Events from nanopore measurements obtained during a 10-min measurement from *Salmonella* Typhimurium. Specificity is shown by selected background events with highlighted additional 16S rRNA ID `3113` events or fragments.

**Fig.S3**

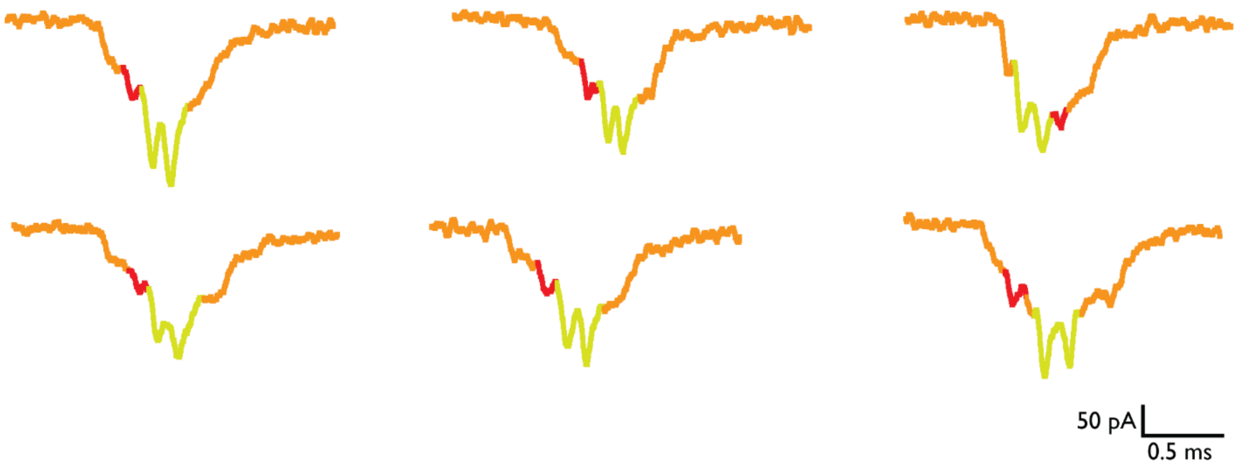


**Fig.S3** Nanopore events for *A. baumannii* 16S rRNA ID `0331`.

**Fig.S4**

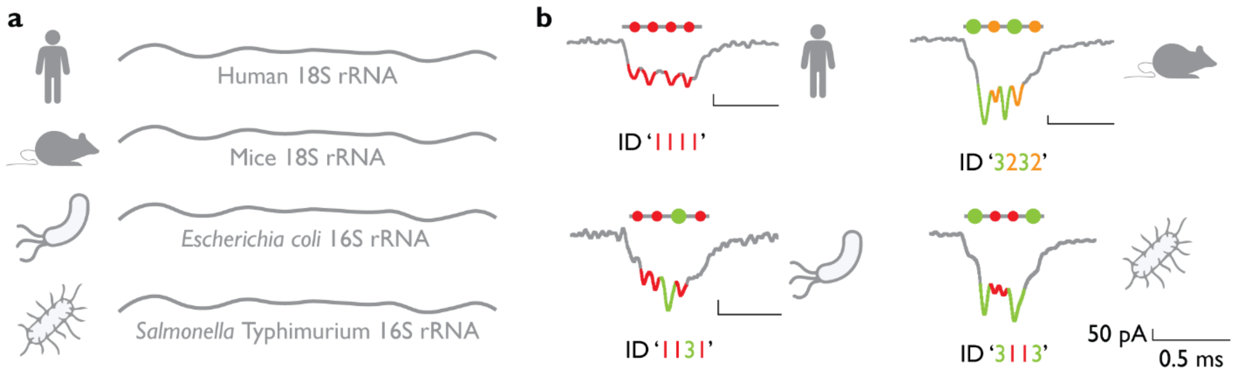


**Fig.S4** Identifiers enable multiplexed species determination using 16S or 18S rRNA and bacterial or viral RNA. (**a**) Total RNA samples from four different sources, humans, mice, E. coli, and S. Typhimurium, were isolated and targeted for either 18S or 16S rRNAs. (**b**) RNA ID designs and their corresponding nanopore readouts are presented to showcase the capability for parallel identification of multiple organisms.

**Fig.S5**

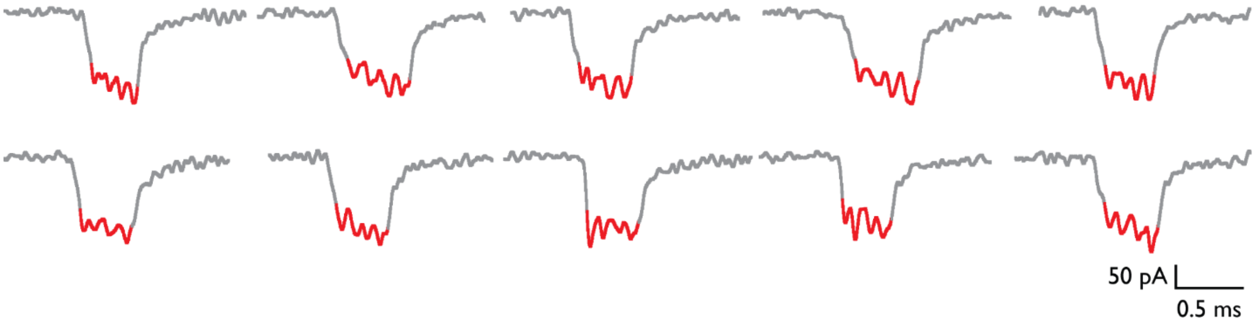


**Fig.S5** Nanopore events for human 18S rRNA ID `1111`.

**Fig.S6**

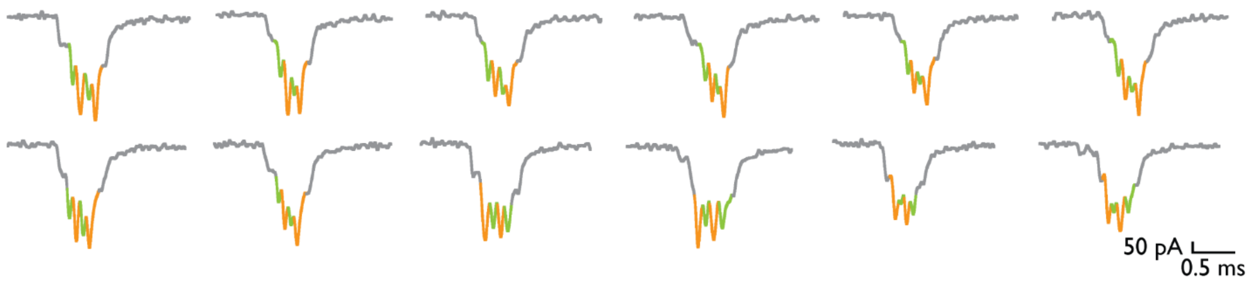


**Fig.S6** Nanopore events for mice 18S rRNA ID `3232`.

**Fig.S7**

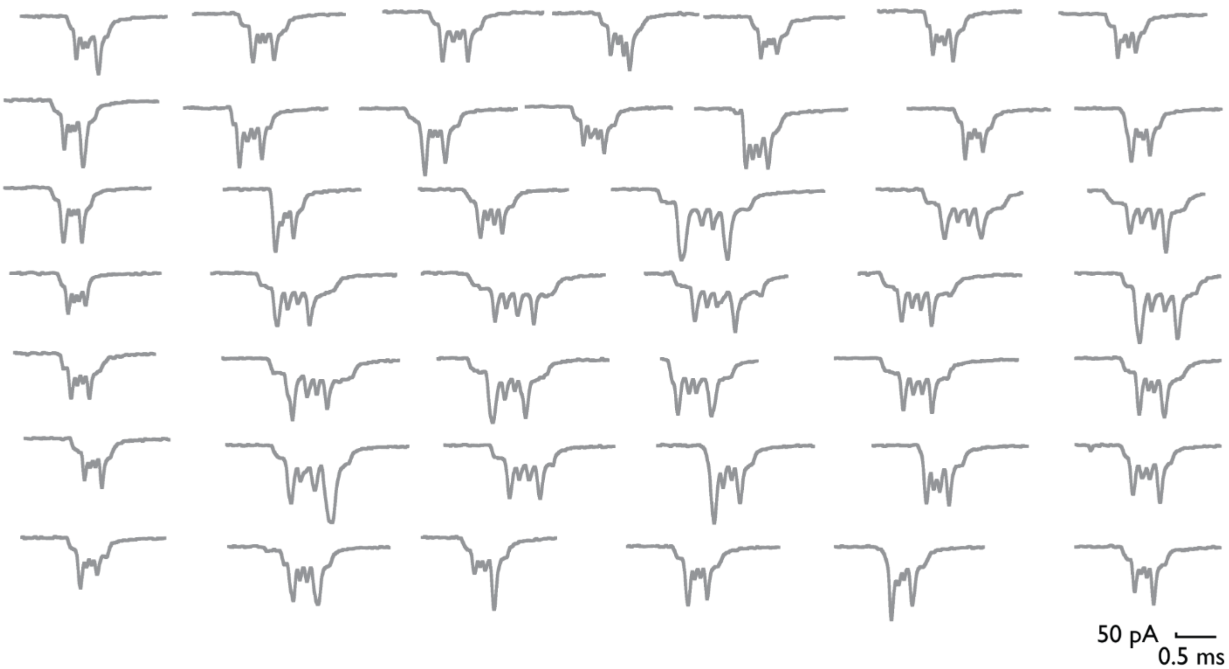


**Fig.S7** Nanopore events for *S.*Typhimurium 16S rRNA ID `3113`.

**Fig.S8**

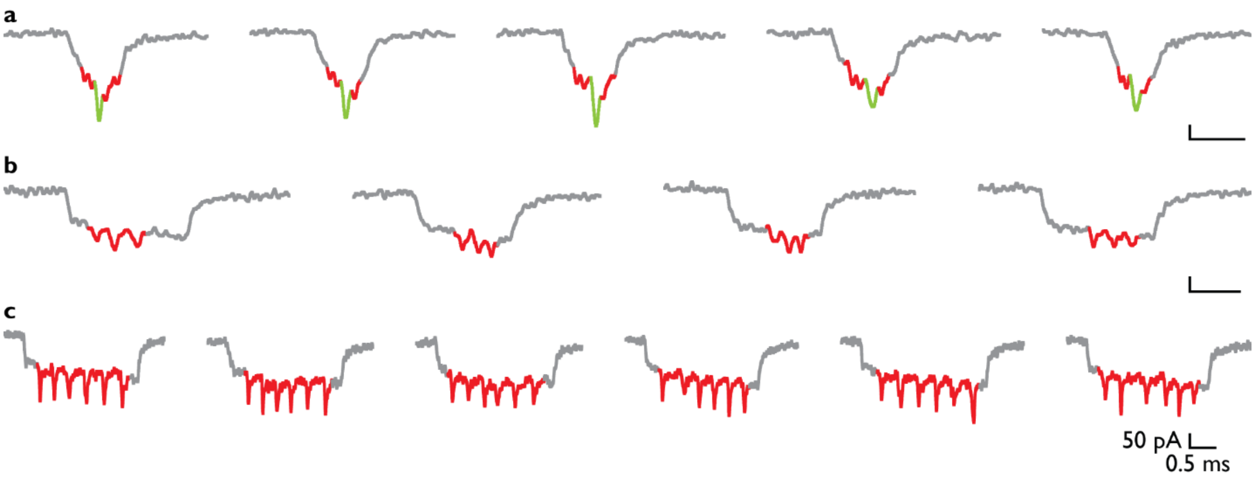


**Fig.S8** Nanopore events for *E.coli* coinfection with single-stranded M13 DNA phage (7,228 nt) and single-stranded MS2 RNA phage (3,569 nt). Besides *E.coli* 16S rRNA ID `1131`, we created M13 ID `111111` and MS2 ID `111`.

**Fig.S9**

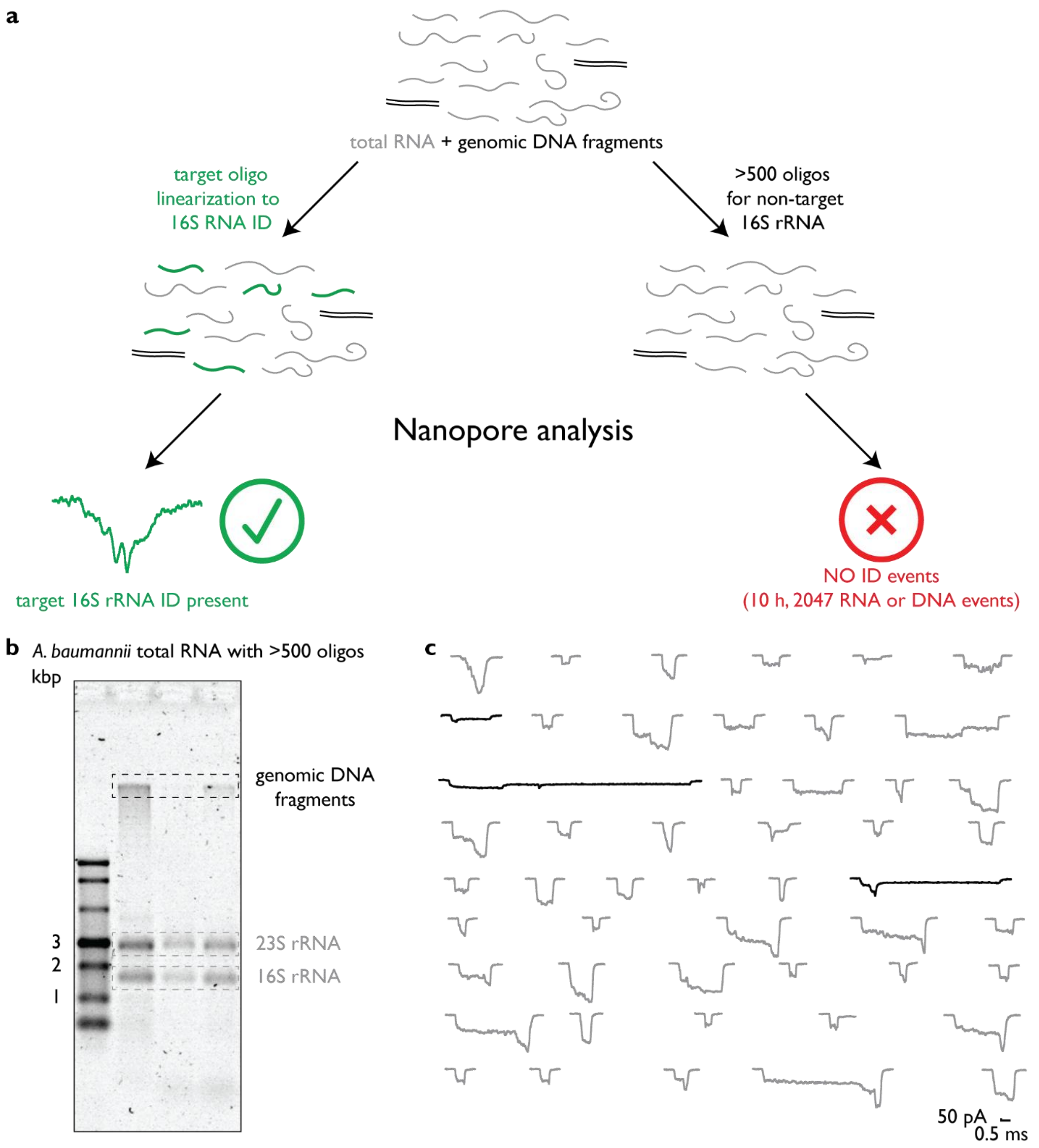


**Fig.S9** Testing for non-specific 16S rRNA ID assembly using total RNA from *A. baumannii*. (**a**) Procedure for assembly RNA IDs and their nanopore analysis is schematically presented. Using the correct oligos targeting 16S rRNA *A. baumannii,* we identified RNA ID \`133\` (as shown in Fig. 4). We tested non-specific RNA ID assembly by mixing almost five hundred oligonucleotides targeting human 18S rRNA, mice 18S rRNA, *E. coli* 16S rRNA, *Salmonella* spp. 16S and 23S

RNA, but by using total RNA from *A. baumannii*. After a 10-h nanopore measurement, we detected 2047 nanopore events originating from background RNA and DNA while none of these had the designed RNA ID. (**b**) 0.8% (v/v) agarose gel electrophoresis shows that there is no shift of 16S rRNA after assembly with >500 oligos. Lane 1 – ssRNA ladder (NEB, catalogue no. N0362S); Lane 2 – total RNA of KL49 *A. baumannii*; Lane 3 – total RNA of KL49 *A. baumannii* assembled with 500 oligos; Lane 4 – total RNA of control *A. baumannii* assembled with 500 oligos. (**c**) 50 consecutive nanopore events when >500 oligos were used with total RNA. RNA blob-like events are shown in grey, and genomic DNA fragments are shown in black. None of these events fulfill requirements for positive RNA ID identification for nanopore events.

**Fig.S10**

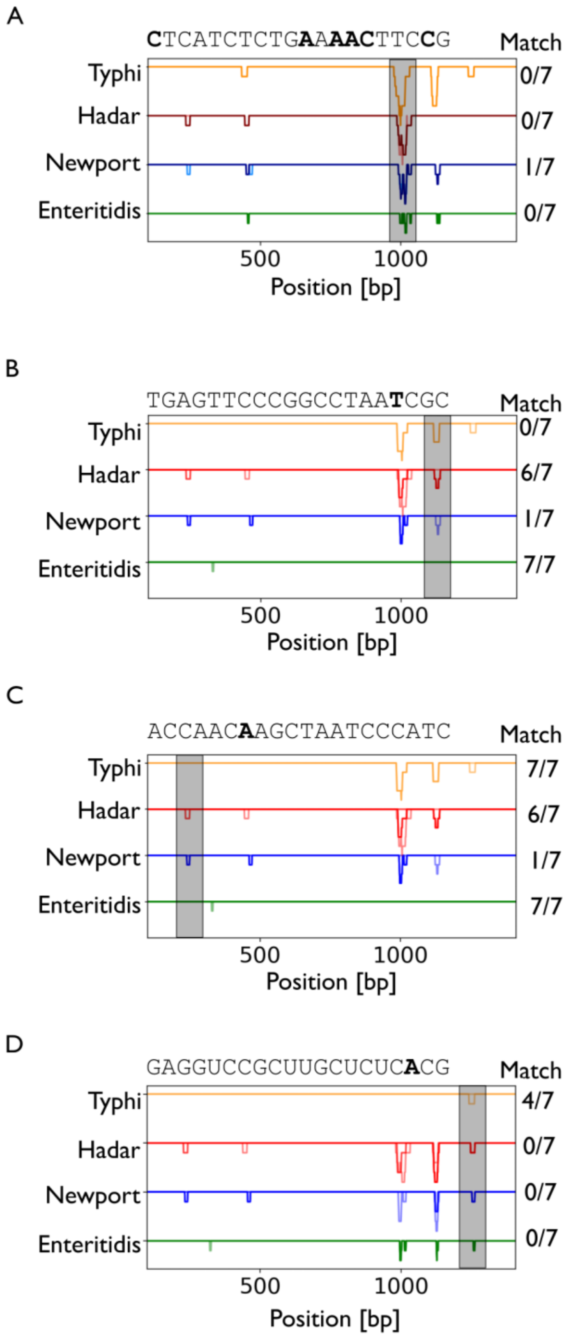


**Fig.S10** Sequence alignments for several locations along rRNA sequences from *Salmonella enterica* serovars. Per bacterium there are seven potential rRNA transcripts originating from seven rRNA genes. Match fraction represents how many sequences the designed guide RNA matches for within the 7 different rRNA sequences.

**Fig.S11**

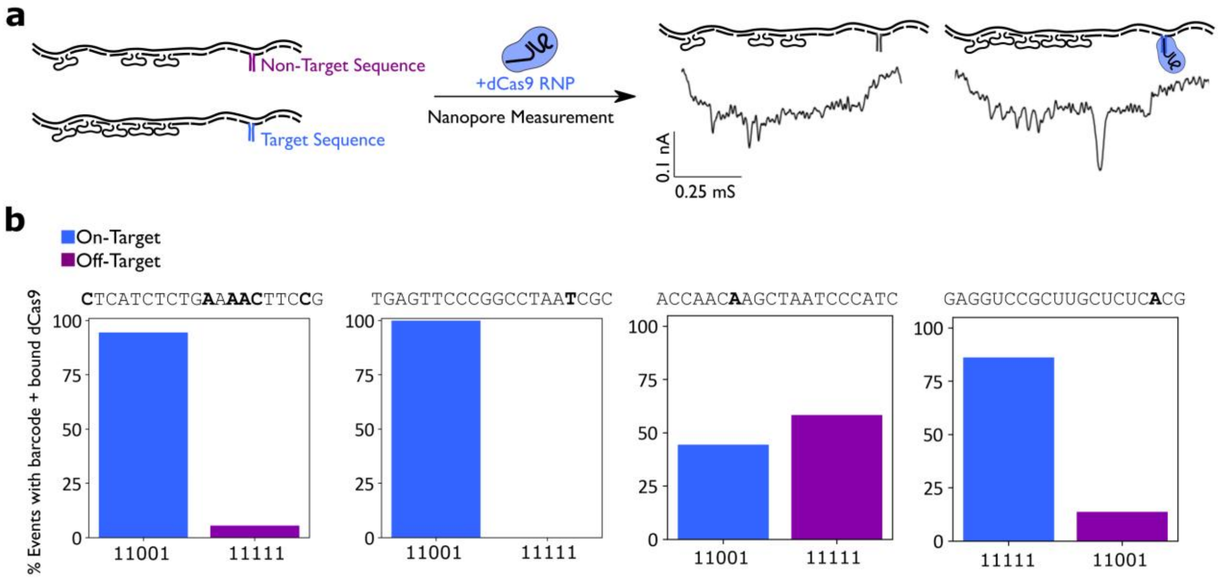


**Fig.S11** Specificity of several probes for target regions to identify single-base pair mismatches. (A) Examples of dCas9 (in blue) bound and unbound nanopore events. (B-D) Specificity of different probes to target sequence developed using method by Sandler et al.[1].

**Fig.S12**

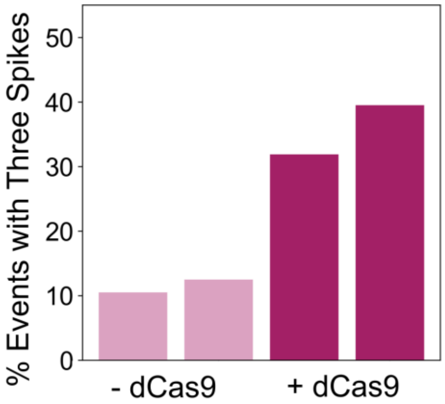


**Fig.S12** Repeats for *S.* Typhi sample. Samples were assembled independently and measured both in the absence (Sample 1, N=25 linear nanopore events, Sample 2, N=16 linear nanopore events) and presence of dCas9 (Sample 1, N=69 linear nanopore events, Sample 2, N=33 linear nanopore events).

**Fig.S13**

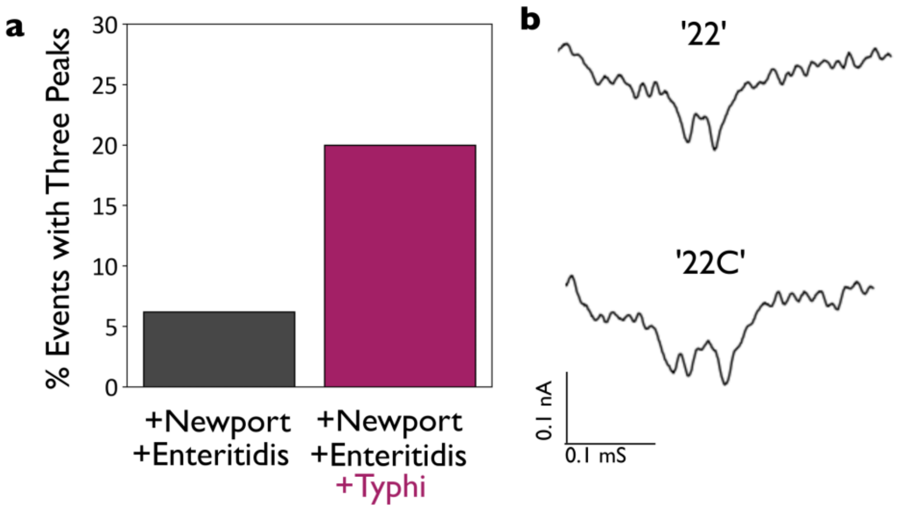


**Fig.S13** *S*. Typhi can be differentiated in samples with mixed strains of *S*. Newport and *S*. Enteritidis using dCas9 by an increase in nanopore events observed with three peaks when *S*. Typhi RNA is mixed in with *S*. Newport and *S*. Enteritidis (**a**). An event with no dCas9 bound (two peaks) is likely *S*. Newport or *S*. Enteritidis (top) and event with dCas9 bound is likely *S*. Typhi (**b**).

### Table S1

**Table S1.** The core DNA oligonucleotides are complementary to linearized ssM13mp18 (7228 nt).

| Oligonucleotide Number | Sequence (5′→3′) | Oligonucleotide Number | Sequence (5′→3′) |
|---|---|---|---|
| 1 | TTTTCGTAATCATGGTCATAGCTGTTTCCTGTGTGAAATTGTTATC | 96 | CTTGAGCCATTTGGGAATTAGAGCCAGCAAAATCACCA |
| 2 | CGCTCACAATTCCACACAACATACGAGCCGGAAGCATA | 97 | GTAGCACCATTACCATTAGCAAGGCCGGAAACGTCACC |
| 3 | AAGTGTAAAGCCTGGGGTGCCTAATGAGTGAGCTAACT | 98 | AATGAAACCATCGATAGCAGCACCGTAATCAGTAGCGA |
| 4 | CACATTAATTGCGTTGCGCTCACTGCCCGCTTTCCAGT | 99 | CAGAATCAAGTTTGCCTTTAGCGTCAGACTGTAGCGCG |
| 5 | CGGGAAACCTGTCGTGCCAGCTGCATTAATGAATCGGC | 100 | TTTTCATCGGCATTTTCGGTCATAGCCCCCTTATTAGC |
| 6 | CAACGCGCGGGGAGAGGCGGTTTGCGTATTGGGCGCCA | 101 | GTTTGCCATCTTTTCATAATCAAAATCACCGGAACCAG |
| 7 | GGGTGGTTTTTCTTTTCACCAGTGAGACGGGCAACAGC | 102 | AGCCACCACCGGAACCGCCTCCCTCAGAGCCGCCACCC |
| 8 | TGATTGCCCTTCACCGCCTGGCCCTGAGAGAGTTGCAG | 103 | TCAGAACCGCCACCCTCAGAGCCACCACCCTCAGAGCC |
| 9 | CAAGCGGTCCACGCTGGTTTGCCCCAGCAGGCGAAAAT | 104 | GCCACCAGAACCACCACCAGAGCCGCCGCCAGCATTGA |
| 10 | CCTGTTTGATGGTGGTTCCGAAATCGGCAAAATCCCTT | 105 | CAGGAGGTTGAGGCAGGTCAGACGATTGGCCTTGATAT |
| 11 | ATAAATCAAAAGAATAGCCCGAGATAGGGTTGAGTGTT | 106 | TCACAAACAAATAAATCCTCATTAAAGCCAGAATGGAA |
| 12 | GTTCCAGTTTGGAACAAGAGTCCACTATTAAAGAACGT | 107 | AGCGCAGTCTCTGAATTTACCGTTCCAGTAAGCGTCAT |
| 13 | GGACTCCAACGTCAAAGGGCGAAAAACCGTCTATCAGG | 108 | ACATGGCTTTTGATGATACAGGAGTGTACTGGTAATAA |
| 14 | GCGATGGCCCACTACGTGAACCATCACCCAAATCAAGT | 109 | GTTTTAACGGGGTCAGTGCCTTGAGTAACAGTGCCCGT |
| 15 | TTTTTGGGGTCGAGGTGCCGTAAAGCACTAAATCGGAA | 110 | ATAAACAGTTAATGCCCCCTGCCTATTTCGGAACCTAT |
| 16 | CCCTAAAGGGAGCCCCCGATTTAGAGCTTGACGGGGAA | 111 | TATTCTGAAACATGAAAGTATTAAGAGGCTGAGACTCC |
| 17 | AGCCGGCGAACGTGGCGAGAAAGGAAGGGAAGAAAGCG | 112 | TCAAGAGAAGGATTAGGATTAGCGGGGTTTTGCTCAGT |
| 18 | AAAGGAGCGGGCGCTAGGGCGCTGGCAAGTGTAGCGGT | 113 | ACCAGGCGGATAAGTGCCGTCGAGAGGGTTGATATAAG |
| 19 | CACGCTGCGCGTAACCACCACACCCGCCGCGCTTAATG | 114 | TATAGCCCGGAATAGGTGTATCACCGTACTCAGGAGGT |
| 20 | CGCCGCTACAGGGCGCGTACTATGGTTGCTTTGACGAG | 115 | TTAGTACCGCCACCCTCAGAACCGCCACCCTCAGAACC |
| 21 | CACGTATAACGTGCTTTCCTCGTTAGAATCAGAGCGGG | 116 | GCCACCCTCAGAGCCACCACCCTCATTTTCAGGGATAG |
| 22 | AGCTAAACAGGAGGCCGATTAAAGGGATTTTAGACAGG | 117 | CAAGCCCAATAGGAACCCATGTACCGTAACACTGAGTT |
| 23 | AACGGTACGCCAGAATCCTGAGAAGTGTTTTTATAATC | 118 | TCGTCACCAGTACAAACTACAACGCCTGTAGCATTCCA |
| 24 | AGTGAGGCCACCGAGTAAAAGAGTCTGTCCATCACGCA | 119 | CAGACAGCCCTCATAGTTAGCGTAACGATCTAAAGTTT |
| 25 | AATTAACCGTTGTAGCAATACTTCTTTGATTAGTAATA | 120 | TGTCGTCTTTCCAGACGTTAGTAAATGAATTTTCTGTA |
| 26 | ACATCACTTGCCTGAGTAGAAGAACTCAAACTATCGGC | 121 | TGGGATTTTGCTAAACAACTTTCAACAGTTTCAGCGGA |
| 27 | CTTGCTGGTAATATCCAGAACAATATTACCGCCAGCCA | 122 | GTGAGAATAGAAAGGAACAACTAAAGGAATTGCGAATA |
| 28 | TTGCAACAGGAAAAACGCTCATGGAAATACCTACATTT | 123 | ATAATTTTTTCACGTTGAAAATCTCCAAAAAAAAGGCT |
| 29 | TGACGCTCAATCGTCTGAAATGGATTATTTACATTGGC | 124 | CCAAAAGGAGCCTTTAATTGTATCGGTTTATCAGCTTG |
| 30 | AGATTCACCAGTCACACGACCAGTAATAAAAGGACAT | 125 | CTTTCGAGGTGAATTTCTTAAACAGCTTGATACCGATA |

| | | | |
|---|---|---|---|
| 31 | TCTGGCCAACAGAGATAGAACCCTTCTGACCTGAAAGC | 126 | GTTGCGCCGACAATGACAACAACCATCGCCCACGCATA |
| 32 | GTAAGAATACGTGGCACAGACAATATTTTTGAATGGCT | 127 | ACCGATATATTCGGTCGCTGAGGCTTGCAGGGAGTTAA |
| 33 | ATTAGTCTTTAATGCGCGAACTGATAGCCCTAAAACAT | 128 | AGGCCGCTTTTGCGGGATCGTCACCCTCAGCAGCGAAA |
| 34 | CGCCATTAAAAATACCGAACGAACCACCAGCAGAAGAT | 129 | GACAGCATCGGAACGAGGGTAGCAACGGCTACAGAGGC |
| 35 | AAAACAGAGGTGAGGCGGTCAGTATTAACACCGCCTGC | 130 | TTTGAGGACTAAAGACTTTTTCATGAGGAAGTTTCCAT |
| 36 | AACAGTGCCACGCTGAGAGCCAGCAGCAAATGAAAAAT | 131 | TAAACGGGTAAAATACGTAATGCCACTACGAAGGCACC |
| 37 | CTAAAGCATCACCTTGCTGAACCTCAAATATCAAACCC | 132 | AACCTAAAACGAAAGAGGCAAAAGAATACACTAAAACA |
| 38 | TCAATCAATATCTGGTCAGTTGGCAAATCAACAGTTGA | 133 | CTCATCTTTGACCCCCAGCGATTATACCAAGCGCGAAA |
| 39 | AAGGAATTGAGGAAGGTTATCTAAAATATCTTTAGGAG | 134 | CAAAGTACAACGGAGATTTGTATCATCGCCTGATAAAT |
| 40 | CACTAACAACTAATAGATTAGAGCCGTCAATAGATAAT | 135 | TGTGTCGAAATCCGCGACCTGCTCCATGTTACTTAGCC |
| 41 | ACATTTGAGGATTTAGAAGTATTAGACTTTACAAACAA | 136 | GGAACGAGGCGCAGACGGTCAATCATAAGGGAACCGAA |
| 42 | TTCGACAACTCGTATTAAATCCTTTGCCCGAACGTTAT | 137 | CTGACCAACTTTGAAAGAGGACAGATGAACGGTGTACA |
| 43 | TAATTTTAAAAGTTTGAGTAACATTATCATTTTGCGGA | 138 | GACCAGGCGCATAGGCTGGCTGACCTTCATCAAGAGTA |
| 44 | ACAAAGAAACCACCAGAAGGAGCGGAATTATCATCATA | 139 | ATCTTGACAAGAACCGGATATTCATTACCCAAATCAAC |
| 45 | TTCCTGATTATCAGATGATGGCAATTCATCAATATAAT | 140 | GTAACAAAGCTGCTCATTCAGTGAATAAGGCTTGCCCT |
| 46 | CCTGATTGTTTGGATTATACTTCTGAATAATGGAAGGG | 141 | GACGAGAAACACCAGAACGAGTAGTAAATTGGGCTTGA |
| 47 | TTAGAACCTACCATATCAAAATTATTTGCACGTAAAAC | 142 | GATGGTTTAATTTCAACTTTAATCATTGTGAATTACCT |
| 48 | AGAAATAAAGAAATTGCGTAGATTTTCAGGTTTAACGT | 143 | TATGCGATTTTAAGAACTGGCTCATTATACCAGTCAGG |
| 49 | CAGATGAATATACAGTAACAGTACCTTTTACATCGGGA | 144 | ACGTTGGGAAGAAAAATCTACGTTAATAAAACGAACTA |
| 50 | GAAACAATAACGGATTCGCCTGATTGCTTTGAATACCA | 145 | ACGGAACAACATTATTACAGGTAGAAAGATTCATCAGT |
| 51 | AGTTACAAAATCGCGCAGAGGCGAATTATTCATTTCAA | 146 | TGAGATTTAGGAATACCACATTCAACTAATGCAGATAC |
| 52 | TTACCTGAGCAAAAGAAGATGATGAAACAAACATCAAG | 147 | ATAACGCCAAAAGGAATTACGAGGCATAGTAAGAGCAA |
| 53 | AAAACAAAATTAATTACATTTAACAATTTCATTTGAAT | 148 | CACTATCATAACCCTCGTTTACCAGACGACGATAAAAA |
| 54 | TACCTTTTTAATGGAAACAGTACATAAATCAATATAT | 149 | CCAAAATAGCGAGAGGCTTTTGCAAAAGAAGTTTTGCC |
| 55 | GTGAGTGAATAACCTTGCTTCTGTAAATCGTCGCTATT | 150 | AGAGGGGGTAATAGTAAAATGTTTAGACTGGATAGCGT |
| 56 | AATTAATTTTCCCTTAGAATCCTTGAAAACATAGCGAT | 151 | CCAATACTGCGGAATCGTCATAAATATTCATTGAATCC |
| 57 | AGCTTAGATTAAGACGCTGAGAAGAGTCAATAGTGAAT | 152 | CCCTCAAATGCTTTAAACAGTTCAGAAAACGAGAATGA |
| 58 | TTATCAAAATCATAGGTCTGAGAGACTACCTTTTTAAC | 153 | CCATAAATCAAAAATCAGGTCTTTACCCTGACTATTAT |
| 59 | CTCCGGCTTAGGTTGGGTTATATAACTATATGTAAATG | 154 | AGTCAGAAGCAAAGCGGATTGCATCAAAAAGATTAAGA |
| 60 | CTGATGCAAATCCAATCGCAAGACAAAGAACGCGAGAA | 155 | GGAAGCCCGAAAGACTTCAAATATCGCGTTTTAATTCG |
| 61 | AACTTTTTCAAATATATTTTAGTTAATTTCATCTTCTG | 156 | AGCTTCAAAGCGAACCAGACCGGAAGCAAACTCCAACA |
| 62 | ACCTAAATTTAATGGTTTGAAATACCGACCGTGTGATA | 157 | GGTCAGGATTAGAGAGTACCTTTAATTGCTCCTTTTGA |
| 63 | AATAAGGCGTTAAATAAGAATAAACACCGGAATCATAA | 158 | TAAGAGGTCATTTTTGCGGATGGCTTAGAGCTTAATTG |
| 64 | TTACTAGAAAAAGCCTGTTTAGTATCATATGCGTTATA | 159 | CTGAATATAATGCTGTAGCTCAACATGTTTTAAATATG |
| 65 | CAAATTCTTACCAGTATAAAGCCAACGCTCAACAGTAG | 160 | CAACTAAAGTACGGTGTCTGGAAGTTTCATTCCATATA |
| 66 | GGCTTAATTGAGAATCGCCATATTTAACAACGCCAACA | 161 | ACAGTTGATTCCCAATTCTGCGAACGAGTAGATTTAGT |

| | | | |
|---|---|---|---|
| 67 | TGTAATTTAGGCAGAGGCATTTTCGAGCCAGTAATAAG | 162 | TTGACCATTAGATACATTTCGCAAATGGTCAATAACCT |
| 68 | AGAATATAAAGTACCGACAAAAGGTAAAGTAATTCTGT | 163 | GTTTAGCTATATTTTCATTTGGGGCGCGAGCTGAAAAG |
| 69 | CCAGACGACGACAATAAACAACATGTTCAGCTAATGCA | 164 | GTGGCATCAATTCTACTAATAGTAGTAGCATTAACATC |
| 70 | GAACGCGCCTGTTTATCAACAATAGATAAGTCCTGAAC | 165 | CAATAAATCATACAGGCAAGGCAAAGAATTAGCAAAAT |
| 71 | AAGAAAAATAATATCCCATCCTAATTTACGAGCATGTA | 166 | TAAGCAATAAAGCCTCAGAGCATAAAGCTAAATCGGTT |
| 72 | GAAACCAATCAATAATCGGCTGTCTTTCCTTATCATTC | 167 | GTACCAAAAACATTATGACCCTGTAATACTTTTGCGGG |
| 73 | CAAGAACGGGTATTAAACCAAGTACCGCACTCATCGAG | 168 | AGAAGCCTTTATTTCAACGCAAGGATAAAAATTTTTAG |
| 74 | AACAAGCAAGCCGTTTTTATTTTCATCGTAGGAATCAT | 169 | AACCCTCATATATTTTAAATGCAATGCCTGAGTAATGT |
| 75 | TACCGCGCCCAATAGCAAGCAAATCAGATATAGAAGGC | 170 | GTAGGTAAAGATTCAAAAGGGTGAGAAAGGCCGGAGAC |
| 76 | TTATCCGGTATTCTAAGAACGCGAGGCGTTTTAGCGAA | 171 | AGTCAAATCACCATCAATATGATATTCAACCGTTCTAG |
| 77 | CCTCCCGACTTGCGGGAGGTTTTGAAGCCTTAAATCAA | 172 | CTGATAAATTAATGCCGGAGAGGGTAGCTATTTTTGAG |
| 78 | GATTAGTTGCTATTTTGCACCCAGCTACAATTTTATCC | 173 | AGATCTACAAAGGCTATCAGGTCATTGCCTGAGAGTCT |
| 79 | TGAATCTTACCAACGCTAACGAGCGTCTTTCCAGAGCC | 174 | GGAGCAAACAAGAGAATCGATGAACGGTAATCGTAAAA |
| 80 | TAATTTGCCAGTTACAAAATAAACAGCCATATTATTTA | 175 | CTAGCATGTCAATCATATGTACCCCGGTTGATAATCAG |
| 81 | TCCCAATCCAAATAAGAAACGATTTTTTGTTTAACGTC | 176 | AAAAGCCCCAAAAACAGGAAGATTGTATAAGCAAATAT |
| 82 | AAAAATGAAAATAGCAGCCTTTACAGAGAGAATAACAT | 177 | TTAAATTGTAAACGTTAATATTTTGTTAAAATTCGCAT |
| 83 | AAAAACAGGGAAGCGCATTAGACGGGAGAATTAACTGA | 178 | TAAATTTTTGTTAAATCAGCTCATTTTTTAACCAATAG |
| 84 | ACACCCTGAACAAAGTCAGAGGGTAATTGAGCGCTAAT | 179 | GAACGCCATCAAAAATAATTCGCGTCTGGCCTTCCTGT |
| 85 | ATCAGAGAGATAACCCACAAGAATTGAGTTAAGCCCAA | 180 | AGCCAGCTTTCATCAACATTAAATGTGAGCGAGTAACA |
| 86 | TAATAAGAGCAAGAAACAATGAAATAGCAATAGCTATC | 181 | ACCCGTCGGATTCTCCGTGGGAACAAACGGCGGATTGA |
| 87 | TTACCGAAGCCCTTTTTAAGAAAAGTAAGCAGATAGCC | 182 | CCGTAATGGGATAGGTCACGTTGGTGTAGATGGGCGCA |
| 88 | GAACAAAGTTACCAGAAGGAAACCGAGGAAACGCAATA | 183 | TCGTAACCGTGCATCTGCCAGTTTGAGGGGACGACGAC |
| 89 | ATAACGGAATACCCAAAAGAACTGGCATGATTAAGACT | 184 | AGTATCGGCCTCAGGAAGATCGCACTCCAGCCAGCTTT |
| 90 | CCTTATTACGCAGTATGTTAGCAAACGTAGAAAATACA | 185 | CCGGCACCGCTTCTGGTGCCGGAAACCAGGCAAAGCGC |
| 91 | TACATAAAGGTGGCAACATATAAAAGAAACGCAAAGAC | 186 | CATTCGCCATTCAGGCTGCGCAACTGTTGGGAAGGGCG |
| 92 | ACCACGGAATAAGTTTATTTTGTCACAATCAATAGAAA | 187 | ATCGGTGCGGGCCTCTTCGCTATTACGCCAGCTGGCGA |
| 93 | ATTCATATGGTTTACCAGCGCCAAAGACAAAAGGGCGA | 188 | AAGGGGGATGTGCTGCAAGGCGATTAAGTTGGGTAACG |
| 94 | CATTCAACCGATTGAGGGAGGGAAGGTAAATATTGACG | 189 | CCAGGGTTTTCCCAGTCACGACGTTGTAAAACGACGGC |
| 95 | GAAATTATTCATTAAAGGTGAATTATCACCGTCACCGA | 190 | CAGTGCCAAGCTTGCATGCCTGCAGGTCGACTCTAGAGGATCTTTT |

**Table S2**

**Table S2.** Barcode 'bits' for DNA nanostructure with ´11111´ and one overhang as shown in Fig. 4a. The following replacements are made to create the ´1´ bits in the barcode portion of the nanostructure.

| **Replaced oligonucleotides from Table S1.** | **Sequence (5′→3′)** |
|---|---|
| **For the first bit:**<br><br>26, 27, 28, 29, 30, 31 and 32 | CTGAAAGCGTAAGAATACGTGGCACAGACAATATTTTTGAATGGCT |
| | ACATCACTTGTCCTCTTTTGAGGAACAAGTTTTCTTGTCCTGAGTAGA |
| | AGAACTCAAATCCTCTTTTGAGGAACAAGTTTTCTTGTCTATCGGCCT |
| | TGCTGGTAATTCCTCTTTTGAGGAACAAGTTTTCTTGTATCCAGAACA |
| | ATATTACCGCTCCTCTTTTGAGGAACAAGTTTTCTTGTCAGCCATTGC |
| | AACAGGAAAATCCTCTTTTGAGGAACAAGTTTTCTTGTACGCTCATGG |
| | AAATACCTACTCCTCTTTTGAGGAACAAGTTTTCTTGTATTTTGACGC |
| | TCAATCGTCTTCCTCTTTTGAGGAACAAGTTTTCTTGTGAAATGGATT |
| | ATTTACATTGTCCTCTTTTGAGGAACAAGTTTTCTTGTGCAGATTCAC |
| | CAGTCACACGTCCTCTTTTGAGGAACAAGTTTTCTTGTACCAGTAATA |
| | AAAGGGACATTCCTCTTTTGAGGAACAAGTTTTCTTGTTCTGGCCAAC |
| | AGAGATAGAATCCTCTTTTGAGGAACAAGTTTTCTTGTCCCTTCTGAC |
| **For the second bit:**<br><br>40, 41, 42, 43, 44, 45 and 46 | AATATAATCCTGATTGTTTGGATTATACTTCTGAATAATGGAAGGG |
| | CACTAACAACTCCTCTTTTGAGGAACAAGTTTTCTTGTTAATAGATTA |
| | GAGCCGTCAATCCTCTTTTGAGGAACAAGTTTTCTTGTTAGATAATAC |
| | ATTTGAGGATTCCTCTTTTGAGGAACAAGTTTTCTTGTTTAGAAGTAT |
| | TAGACTTTACTCCTCTTTTGAGGAACAAGTTTTCTTGTAAACAATTCG |
| | ACAACTCGTATCCTCTTTTGAGGAACAAGTTTTCTTGTTTAAATCCTT |
| | TGCCCGAACGTCCTCTTTTGAGGAACAAGTTTTCTTGTTTATTAATTT |
| | TAAAAGTTTGTCCTCTTTTGAGGAACAAGTTTTCTTGTAGTAACATTA |
| | TCATTTTGCGTCCTCTTTTGAGGAACAAGTTTTCTTGTGAACAAAGAA |
| | ACCACCAGAATCCTCTTTTGAGGAACAAGTTTTCTTGTGGAGCGGAAT |
| | TATCATCATATCCTCTTTTGAGGAACAAGTTTTCTTGTTTCCTGATTA |
| | TCAGATGATGTCCTCTTTTGAGGAACAAGTTTTCTTGTGCAATTCATC |
| **For the third bit:** | TGTAAATGCTGATGCAAATCCAATCGCAAGACAAAGAACGCGAGAA |

| | |
|---|---|
| 54, 55, 56, 57, 58, 59 and 60 | TACCTTTTTTTCCTCTTTTGAGGAACAAGTTTTCTTGTAATGGAAACA |
| | GTACATAAATTCCTCTTTTGAGGAACAAGTTTTCTTGTCAATATATGT |
| | GAGTGAATAATCCTCTTTTGAGGAACAAGTTTTCTTGTCCTTGCTTCT |
| | GTAAATCGTCTCCTCTTTTGAGGAACAAGTTTTCTTGTGCTATTAATT |
| | AATTTTCCCTTCCTCTTTTGAGGAACAAGTTTTCTTGTTAGAATCCTT |
| | GAAAACATAGTCCTCTTTTGAGGAACAAGTTTTCTTGTCGATAGCTTA |
| | GATTAAGACGTCCTCTTTTGAGGAACAAGTTTTCTTGTCTGAGAAGAG |
| | TCAATAGTGATCCTCTTTTGAGGAACAAGTTTTCTTGTATTTATCAAA |
| | ATCATAGGTCTCCTCTTTTGAGGAACAAGTTTTCTTGTTGAGAGACTA |
| | CCTTTTTAACTCCTCTTTTGAGGAACAAGTTTTCTTGTCTCCGGCTTA |
| | GGTTGGGTTATCCTCTTTTGAGGAACAAGTTTTCTTGTTATAACTATA |
| **For the fourth bit:** | TCATCGAGAACAAGCAAGCCGTTTTTATTTTCATCGTAGGAATCAT |
| 68, 69, 70, 71, 72, 73 and 74 | AGAATATAAATCCTCTTTTGAGGAACAAGTTTTCTTGTGTACCGACAA |
| | AAGGTAAAGTTCCTCTTTTGAGGAACAAGTTTTCTTGTAATTCTGTCC |
| | AGACGACGACTCCTCTTTTGAGGAACAAGTTTTCTTGTAATAAACAAC |
| | ATGTTCAGCTTCCTCTTTTGAGGAACAAGTTTTCTTGTAATGCAGAAC |
| | GCGCCTGTTTTCCTCTTTTGAGGAACAAGTTTTCTTGTATCAACAATA |
| | GATAAGTCCTTCCTCTTTTGAGGAACAAGTTTTCTTGTGAACAAGAAA |
| | AATAATATCCTCCTCTTTTGAGGAACAAGTTTTCTTGTCATCCTAATT |
| | TACGAGCATGTCCTCTTTTGAGGAACAAGTTTTCTTGTTAGAAACCAA |
| | TCAATAATCGTCCTCTTTTGAGGAACAAGTTTTCTTGTGCTGTCTTTC |
| | CTTATCATTCTCCTCTTTTGAGGAACAAGTTTTCTTGTCAAGAACGGG |
| | TATTAAACCATCCTCTTTTGAGGAACAAGTTTTCTTGTAGTACCGCAC |
| **For the fifth bit:** | AGATAGCCGAACAAAGTTACCAGAAGGAAACCGAGGAAACGCAATA |
| 82, 83, 84, 85, 86, 87 and 88 | AAAAATGAAATCCTCTTTTGAGGAACAAGTTTTCTTGTATAGCAGCCT |
| | TTACAGAGAGTCCTCTTTTGAGGAACAAGTTTTCTTGTAATAACATAA |
| | AAACAGGGAATCCTCTTTTGAGGAACAAGTTTTCTTGTGCGCATTAGA |
| | CGGGAGAATTTCCTCTTTTGAGGAACAAGTTTTCTTGTAACTGAACAC |
| | CCTGAACAAATCCTCTTTTGAGGAACAAGTTTTCTTGTGTCAGAGGGT |
| | AATTGAGCGCTCCTCTTTTGAGGAACAAGTTTTCTTGTTAATATCAGA |
| | GAGATAACCCTCCTCTTTTGAGGAACAAGTTTTCTTGTACAAGAATTG |

| |
|---|
| AGTTAAGCCCTCCTCTTTTGAGGAACAAGTTTTCTTGTAATAATAAGA |
| GCAAGAAACATCCTCTTTTGAGGAACAAGTTTTCTTGTATGAAATAGC |
| AATAGCTATCTCCTCTTTTGAGGAACAAGTTTTCTTGTTTACCGAAGC |
| CCTTTTTAAGTCCTCTTTTGAGGAACAAGTTTTCTTGTAAAAGTAAGC |

**Table S3**

**Table S3.** Overhang sequences for DNA nanostructure with ´11111´ and one overhang as shown in Fig.4a. The sequences for overhangs are labelled as: Bold – crRNA target region, Red – Mutation, Blue – PAM).

| Serovar_replaced oligonucleotides from Table S1. | Sequence (5′→3′) |
|---|---|
| Typhi_142 | GATGGTTTAATTTCAACTTTAATCATTGTGAATTACCTTTCAAAGAGAA GCGACCTCGTGAGAGCAAGCGGACCTCATAAAGTGCGTCGT |
| Typhi_143 | ACGACGCACTTTAT**GAGGTCCGCTTGCTCTCACGAGG**TCGCTTCTCTTT GTTTATGCGATTTTAAGAACTGGCTCATTATACCAGTCAGG |
| Newport_142 | GATGGTTTAATTTCAACTTTAATCATTGTGAATTACCTTTCAAAGAGAA GCGACCTCGCGAGAGCAAGCGGACCTCATAAAGTGCGTCGT |
| Newport_143 | ACGACGCACTTTAT**GAGGTCCGCTTGCTCTCGCGAGG**TCGCTTCTCTTT GTTTATGCGATTTTAAGAACTGGCTCATTATACCAGTCAGG |

**Table S4**

**Table S4.** Oligonucleotides to assemble RNA ID 'C22' designed for serovar *Salmonella* Typhi. Lowercase portions are overhangs for biotinylated sequences to bind to and bold is the target area of the dCas9.

| Oligonucleotide Number | Sequence (5′→3′) |
|---|---|
| 1 | AAGGAGGTGATCCAACCGCAGGTTCCCCTACGGTTACCTT |
| 2 | GTTACGACTTCACCCCAGTCATGAATCACAAAGTGGTAAG |
| 3 | CGCCCTCCCGAAGGTTAAGCTACCTACTTCTTTTGCAACC |
| 4 | CACTCCCATGGTGTGACGGGCGGTGTGTACAAGGCCCGGG |
| 5 | AACGTATTCACCGTGGCATTCTGATCCACGATTACTAGCG |
| 6 | ATTCCGACTTCATGGAGTCGAGTTGCAGACTCCAATCCGGACTAC |
| 7 | GACGCACTTTAT**GAGGTCCGCTTGCTCTCACG**AGGTCGCT |
| 8 | TCTCTTTGTATGCGCCATTGTAGCACGTGTGTAGC |
| 9 | CCTGGTCGTAAGGGCCATGATGACTTGACGTCATCCCCAC |
| 10 | CTTCCTCCAGTTTATCACTGGCAGTCTCCTTTGAGTTCCC |
| 11 | GGCCGGACCGCTGGCAACAAAGGATAAGGGTTGCGCTCGT |
| 12 | TGCGGGACTTAACCCAACATTTCACAACACGAGCTGACGA |
| 13 | CAGCCATGCAGCACCTGTCTCACAGTTCCCGAAGGCACCA |
| 14 | ATCCATCTCTGGAAAGTTCTGTGGATGTCAAGACCAGGTA |
| 15 | AGGTTCTTCGCGTTGCATCGAATTAAACCACATGCTCCAC |
| 16 | CGCTTGTGCGGGCCCCCGTCAATTCATTTGAGTTTTAACC |
| 17 | TTGCGGCCGTACTCCCCAGGCGGTCTACTTAACGCGTTAG |
| 18 | CTCCGGAAGCCACGCCTCAAGGGCACAACCTCCAAGTAGA |
| 19 | CATCGTTTACGGCGTGGACTACCAGGGTATCTAATCCTGT |
| 20 | TTGCTCCCCACGCTTTCGCACCTGAGCGTCAGTCTTTGTC |
| 21 | CAGGGGGCCGCCTTCGCCACCGGTATTCCTCCAGATCTCT |
| 22 | ACGCATTTCACCGCTACACCTGGAATTCTACCCCCCTCTA |
| 23 | CAAGACTCAAGCCTGCCAGTTTCGAATGCAGTTCCCAGGT |
| 24 | TGAGCCCGGGGATTTCACATCCGACTTGACAGACCGCCTG |
| 25 | CGTGCGCTTTACGCCCAGTAATTC |
| 26 | CGATTAACGCTTGCACCCTCCGTA |
| 27 | TTACCGCGGCTGCTGGCACGGAGTTAGCCGtttggatatcactcattagtggt |
| 28 | GTGCTTCTTCTGCGGGTAACGTCAATTGCTtttggatatcactcattagtggt |
| 29 | GCGGTTATTAACCACAACACCTTCCTCCCCGCTGAAAGTA |
| 30 | CTTTACAACCCGAAGGCCTTCTTCATACACGCGGCATGGC |
| 31 | TGCATCAGGCTTGCGCCCATTGTGCAATATTCCCCACTGC |
| 32 | TGCCTCCCGTAGGAGTCTGGACCGT |
| 33 | GTCTCAGTTCCAGTGTGGCTGGTCA |
| 34 | TCCTCTCAGACCAGCTAGGGATCGTCGCCTtttggatatcactcattagtggt |

| | |
|---|---|
| 35 | TGGTGAGCCGTTACCTCACCAACAAGCTAAtttggatatcactcattagtggt |
| 36 | TCCCATCTGGGCACATCTGATGGCAAGAGGCCCGAAGGTC |
| 37 | CCCCTCTTTGGTCTTGCGACGTTATGCGGTATTAGCCACC |
| 38 | GTTTCCAGTAGTTATCCCCCTCCATCAGGCAGTTTCCCAG |
| 39 | ACATTACTCACCCGTCCGCCACTCGTCAGCAAAGCAGCA |
| 40 | AGCTGCTTCCTGTTACCGTTCGACTTGCATGTGTTAGGCC |
| 41 | TGCCGCCAGCGTTCAATCTGAGCCATGATCAAACTCT |

### Table S5

**Table S5.** *E.coli* J01859.1 16S rRNA oligonucleotide sequences for ID `1131`.

| Oligonucleotide Number | Sequence (5′→3′) |
|---|---|
| 1 | TAAGGAGGTGATCCAACCGCAGGTTCCCCTACGGTTAC |
| 2 | CTTGTTACGACTTCACCCCAGTCATGAATCACAAAGTG |
| 3 | GTAAGCGCCCTCCCGAAGGTTAAGCTACCTACTTCTTT |
| 4 | TGCAACCCACTCCCATGGTGTGACGGGCGGTGTGTACA |
| 5 | AGGCCCGGGAACGTATTCACCGTGGCATTCTGATCCAC |
| 6 | GATTACTAGCGATTCCGACTTCATGGAGTCGAGTTGCA |
| 7 | GACTCCAATCCGGACTACGACGCACTTTATGAGGTCCG |
| 8 | CTTGCTCTCGCGAGGTCGCTTCTCTTTGTATGCGCCAT |
| 9 | TGTAGCACGTGTGTAGCCCTGGTCGTAAGGGCCATGAT |
| 10 | GACTTGACGTCATCCCCACCTTCCTCCAGTTTATCACT |
| 11 | GGCAGTCTCCTTTGAGTT |
| 12 | CCCGGCCGGACCGCTGGCAACAAAGtttggatatcactcattagtggt |
| 13 | GATAAGGGTTGCGCTCGTTGCGGGACTTAACCCAACAT |
| 14 | TTCACAACACGAGCTGACGACAGCCATGCAGCACCTGT |
| 15 | CTCACGGTTCCCGAAGGCACATTCTCATCTCTGAAAAC |
| 16 | TTCCGTGGATGTCAAGACCAGGTAAGGTTCTTCGCGTT |
| 17 | GCATCGAATTAAACCACATGCTC |
| 18 | CACCGCTTGTGCGGGCCCCCGTCAAtttggatatcactcattagtggt |
| 19 | TTCATTTGAGTTTTAACCTTGCGGCCGTACTCCCCAGG |
| 20 | CGGTCGACTTAACGCGTTAGCTCCGGAAGCCACGCCTC |
| 21 | AAGGGCACAACCTCCAAGTCGACATCGTTTACGGCGTG |
| 22 | GACTACCAGGGTATCTAATCCTGTTTGCTCCCCACGCT |
| 23 | TTCGCACCTGAGCGTCAGTCTTC |
| 24 | GTCCAGGGGGCCGCCTTCGCCACCGtttggatatcactcattagtggt |
| 25 | GTATTCCTCCAGATCTCTACGCATTtttggatatcactcattagtggt |
| 26 | TCACCGCTACACCTGGAATTCTACCtttggatatcactcattagtggt |
| 27 | CCCCTCTACGAGACTCAAGCTTGCCAGTATCAGATGCA |
| 28 | GTTCCCAGGTTGAGCCCGGGGATTTCACATCTGACTTA |
| 29 | ACAAACCGCCTGCGTGCGCTTTACGCCCAGTAATTCCG |
| 30 | ATTAACGCTTGCACCCTCCGTATTAC |
| 31 | CGCGGCTGCTGGCACGGAGTTAGCC |
| 32 | GGTGCTTCTTCTGCGGGTAACGTCAtttggatatcactcattagtggt |
| 33 | ATGAGCAAAGGTATTAACTTTACTCCCTTCCTCCCCGC |
| 34 | TGAAAGTACTTTACAACCCGAAGGCCTTCTTCATACAC |
| 35 | GCGGCATGGCTGCATCAGGCTTGCGCCCATTGTGCAAT |
| 36 | ATTCCCCACTGCTGCCTCCCGTAGGAGTCTGGACCGTG |

| | |
|---|---|
| 37 | TCTCAGTTCCAGTGTGGCTGGTCATCCTCTCAGACCAG |
| 38 | CTAGGGATCGTCGCCTAGGTGAGCCGTTACCCCACCTA |
| 39 | CTAGCTAATCCCATCTGGGCACATCCGATGGCAAGAGG |
| 40 | CCCGAAGGTCCCCCTCTTTGGTCTTGCGACGTTATGCG |
| 41 | GTATTAGCTACCGTTTCCAGTAGTTATCCCCCTCCATC |
| 42 | AGGCAGTTTCCCAGACATTACTCACCCGTCCGCCACTC |
| 43 | GTCAGCAAAGAGCAAGCTTCTTCCTGTTACCGTTCGAC |
| 44 | TTGCATGTGTTAGGCCTGCCGCCAGCGTTCAATCTGAG |
| 45 | CCATGATCAAACTCTTCAATTT |

### Table S6

**Table S6.** *A. baumannii* 16S rRNA oligonucleotide sequences for ID `0331`.

| Oligonucleotide Number | Sequence (5′→3′) |
|---|---|
| 1 | TAAGGAGGTGATCCAGCCGCAGGTTCCCCTACGGCTAC |
| 2 | CTTGTTACGACTTCACCCCAGTCATCGGCCACACCGTG |
| 3 | GTAACCGCCCTCTTTGCAGTTAGGCTAGCTACTTCTGG |
| 4 | TGCAACAAACTCCCATGGTGTGACGGGCGGTGTGTACA |
| 5 | AGGCCCGGGAACGTATTCACCGCGGCATTCTGATCCGC |
| 6 | GATTACTAGCGATTCCGACTTCATGGAGTCGAGTTGCA |
| 7 | GACTCCAATCCGGACTACGATCGGCTTTTTGAGATTAG |
| 8 | CATCACATCGCTGTGTAGCAACCCTTTGTACCGACCAT |
| 9 | TGTAGCACGTGTGTAGCCCTGGCCGTAAGGGCCATGAT |
| 10 | GACTTGACGTCGTCCCCGCCTTCCTCCAGTTTGTCACT |
| 11 | GGCAGTATCCTTAAAGTTCCCATCCGAAATGCTGGCAA |
| 12 | GTAAGGAAAAGGGTTGCGCTCGTTGCGGGACTTAACCC |
| 13 | AACATCTCACGACACGAGCTGACGACAGCCATGCAGCA |
| 14 | CCTGTATCTAGATTCCCGAAGGCACCAATCCATCTCTG |
| 15 | GAAAGTTTCTAGTATGTCAAGGCCAGGTAAGGTTCTTC |
| 16 | GCGTTGCATCGAATTAAACCACAT |
| 17 | GCTCCACCGCTTGTGCGGGCCCCCGtttggatatcactcattagtggt |
| 18 | TCAATTCATTTGAGTTTTAGTCTTGtttggatatcactcattagtggt |
| 19 | CGACCGTACTCCCCAGGCGGTCTACtttggatatcactcattagtggt |
| 20 | TTATCGCGTTAGCTGCGCCACTAAAGCCTCAAAGGCCC |
| 21 | CAACGGCTAGTAGACATCGTTTACGGCATGGACTACCA |
| 22 | GGGTATCTAATCCTGTTTGCTCCCCATGCTTTCGTACC |
| 23 | TCAGCGTCAGTATTAGGCCAGATGGCTGCCTTCGCCAT |
| 24 | CGGTATTCCTCCAGATCTCTACG |
| 25 | CATTTCACCGCTACACCTGGAATTCtttggatatcactcattagtggt |
| 26 | TACCATCCTCTCCCATACTCTAGCTtttggatatcactcattagtggt |
| 27 | CACCAGTATCGAATGCAATTCCCAAtttggatatcactcattagtggt |
| 28 | GTTAAGCTCGGGGATTTCACATCCGACTTAATAAGCCG |
| 29 | CCTACGCACGCTTTACGCCCAGTAAATCCGATTAACGC |
| 30 | TCGCACCCTCTGTATTACCGCGGCTGCTGGCACAGAGT |
| 31 | TAGCCGGTGCTTATTCTGCGAGTAACGTCCACTATCTC |
| 32 | TAGGTATTAACTAAAGTAGCCTC |
| 33 | CTCCTCGCTTAAAGTGCTTTACAACtttggatatcactcattagtggt |
| 34 | CATAAGGCCTTCTTCACACACGCGGCATGGCTGGATCA |
| 35 | GGGTTCCCCCCATTGTCCAATATTCCCCACTGCTGCCT |
| 36 | CCCGTAGGAGTCTGGGCCGTGTCTCAGTCCCAGTGTGG |

| | |
|---|---|
| 37 | CGGATCATCCTCTCAGACCCGCTACAGATCGTCGCCTT |
| 38 | GGTAGGCCTTTACCCCACCAACTAGCTAATCCGACTTA |
| 39 | GGCTCATCTATTAGCGCAAGGTCCGAAGATCCCCTGCT |
| 40 | TTCTCCCGTAGGACGTATGCGGTATTAGCATCCCTTTC |
| 41 | GAGATGTTGTCCCCCACTAATAGGCAGATTCCTAAGCA |
| 42 | TTACTCACCCGTCCGCCGCTAGGTCCAGTAGCAAGCTA |
| 43 | CCTTCCCCCGCTCGACTTGCATGTGTTAAGCCTGCCGC |
| 44 | CAGCGTTCAATCTGAGCCATGATCAAACTCTTCAGTTA |

### Table S7

**Table S7.** *A. baumannii* gp100 mRNA oligonucleotide sequences for ID `3113`.

| Oligonucleotide Number | Sequence (5′→3′) |
|---|---|
| 1 | TTACGCATTATTAACTAAATTCCATATCTTATGAAAAT |
| 2 | CTGTATTTTGATTCCACCATAATACTTTAAAACCCATT |
| 3 | GCTTCATATTCTTTACCCAAAGTAGACTCATTACATAA |
| 4 | GCAAACGTTAGGATATGGAAATCTTTT |
| 5 | TCCTAAACATAATTCATCATTTTCAAC |
| 6 | TACACTTTTTATTTCTTCATTAATAtttggatatcactcattagtggt |
| 7 | TAAAGTGGATGAGGTCTAAAAAAAAtttggatatcactcattagtggt |
| 8 | CTTTAATCCCTTTATCTTTCAAGAAtttggatatcactcattagtggt |
| 9 | TTTATATAATAACAGTTCCTGATTTATAAAAAATGGAT |
| 10 | TACTAATTATCAATACTTCACATTTATCAGAAAAGTTG |
| 11 | ATATCTAAACTAATTTTCATATAATCAATTTTATTAGG |
| 12 | TATACCCTCTAATATATTTTTTAGAAAAATACTCAATT |
| 13 | GAGATTTATCAAAAACTTTTAAA |
| 14 | AAAGAAATATTTGGTATTTTATGCTtttggatatcactcattagtggt |
| 15 | TAACAACATAATTTTCTGTTAAAATTCCATGCTGTATT |
| 16 | ATCTCAATATCAAAATTACCAATATTTGACAAAACAGT |
| 17 | TAACCACCTATCATAATGATTTACCACACATATTTTAT |
| 18 | TTATATTTCTTTTATTTAATACAATAACAAATTCATAA |
| 19 | AATAAAATTAATTCAAAAATTGC |
| 20 | TGTTAAGTGTAAATAATATTTAATAtttggatatcactcattagtggt |
| 21 | TTAACAAAAAATATCATTTTAAAGTAAAGTATAATTGA |
| 22 | ATGAATAAAACTCTTCAGGATAACTAGAGGACTTAAAT |
| 23 | AGCCAACAAAACAATAAACTTTATTATTAATACATTGC |
| 24 | AAAAACTCTATATCAGAAGCATATTCTTCATAAATTGC |
| 25 | AATTACTTTTGGAGAGTTAGAAA |
| 26 | AAGATAAAAAATATTGATCATTTTCtttggatatcactcattagtggt |
| 27 | AAGATTTGATGTATTTCTACTCCTTtttggatatcactcattagtggt |
| 28 | CCAAAAAAAACAAAAAATTCAATAGtttggatatcactcattagtggt |
| 29 | CAAAATAAATTGGTATTAAAATAGAAGATAGCACAGAA |
| 30 | ATCCATAAAGGAATTTTATCTTTGAATCGTAGCATTTT |
| 31 | GAAAATAGATAGCTTTTTCGAGCCATAATTTAGCTTCT |
| 32 | CAAAAAGATAATTATCTACTGAAATTTCATTTTCAATA |
| 33 | GCTTTTTTTATAATTTCATA |
| 34 | TTTCTTATTTTTTAGGCACAT |

### Table S8

**Table S8.** Human 18S rRNA oligonucleotide sequences for ID `1111`.

| Oligonucleotide Number | Sequence (5′→3′) |
|---|---|
| 1 | TAATGATCCTTCCGCAGGTTCACCTACGGAAACCTTGT |
| 2 | TACGACTTTTACTTCCTCTAGATAGTCAAGTTCGACCG |
| 3 | TCTTCTCAGCGCTCCGCCAGGGCCGTGGGCCGACCCCG |
| 4 | GCGGGGCCGATCCGAGGGCCTCACTAAACCATCCAATC |
| 5 | GGTAGTAGCGACGGGCGGTGTGTACAAAGGGCAGGGAC |
| 6 | TTAATCAACGCAAGCTTATGACCCGCACTTACTGGGAA |
| 7 | TTCCTCGTTCATGGGGAATAATTGC |
| 8 | AATCCCCGATCCCCATCACGAATGG |
| 9 | GGTTCAACGGTCCTCTTTTGAGGAACAAGTTTTCTTGTGTTACCCGCG |
| 10 | CCTGCCGGCGTCCTCTTTTGAGGAACAAGTTTTCTTGTTAGGGTAGGC |
| 11 | ACACGCTGAGTCCTCTTTTGAGGAACAAGTTTTCTTGTCCAGTCAGTG |
| 12 | TAGCGCGCGTTCCTCTTTTGAGGAACAAGTTTTCTTGTGCAGCCCCGG |
| 13 | ACATCTAAGGTCCTCTTTTGAGGAACAAGTTTTCTTGTGCATCACAGA |
| 14 | CCTGTTATTGTCCTCTTTTGAGGAACAAGTTTTCTTGTCTCAATCTCG |
| 15 | GGTGGCTGAACGCCACTTGTCCCTCTAAGAAGTTGGGG |
| 16 | GACGCCGACCGCTCGGGGGTCGCGTAACTAGTTAGCAT |
| 17 | GCCAGAGTCTCGTTCGTTATCGGAATTAACCAGACAAA |
| 18 | TCGCTCCACCAACTAAGAACGGCCATGCACCACCACCC |
| 19 | ACGGAATCGAGAAAGAGCTATCAATCTGTCAATCCTGT |
| 20 | CCGTGTCCGGGCCGGGTGAGGTTTCCCGTGTTGAGTCA |
| 21 | AATTAAGCCGCAGGCTCCACTCCTG |
| 22 | GTGGTGCCCTTCCGTCAATTCCTTT |
| 23 | AAGTTTCAGCTCCTCTTTTGAGGAACAAGTTTTCTTGTTTTGCAACCA |
| 24 | TACTCCCCCCTCCTCTTTTGAGGAACAAGTTTTCTTGTGGAACCCAAA |
| 25 | GACTTTGGTTTCCTCTTTTGAGGAACAAGTTTTCTTGTTCCCGGAAGC |
| 26 | TGCCCGGCGGTCCTCTTTTGAGGAACAAGTTTTCTTGTGTCATGGGAA |
| 27 | TAACGCCGCCTCCTCTTTTGAGGAACAAGTTTTCTTGTGCATCGCCGG |
| 28 | TCGGCATCGTTCCTCTTTTGAGGAACAAGTTTTCTTGTTTATGGTCGG |
| 29 | AACTACGACGGTATCTGATCGTCTTCGAACCTCCGACT |
| 30 | TTCGTTCTTGATTAATGAAAACATTCTTGGCAAATGCT |
| 31 | TTCGCTCTGGTCCGTCTTGCGCCGGTCCAAGAATTTCA |
| 32 | CCTCTAGCGGCGCAATACGAATGCCCCCGGCCGTCCCT |
| 33 | CTTAATCATGGCCTCAGTTCCGAAAACCAACAAAATAG |
| 34 | AACCGCGGTCCTATTCCATTATTCCTAGCTGCGGTATC |
| 35 | CAGGCGGCTCGGGCCTGCTTTGAAC |
| 36 | ACTCTAATTTTTTCAAAGTAAACGC |

| | |
|---|---|
| 37 | TTCGGGCCCCTCCTCTTTTGAGGAACAAGTTTTCTTGTGCGGGACACT |
| 38 | CAGCTAAGAGTCCTCTTTTGAGGAACAAGTTTTCTTGTCATCGAGGGG |
| 39 | GCGCCGAGAGTCCTCTTTTGAGGAACAAGTTTTCTTGTGCAAGGGGCG |
| 40 | GGGACGGGCGTCCTCTTTTGAGGAACAAGTTTTCTTGTGTGGCTCGCC |
| 41 | TCGCGGCGGATCCTCTTTTGAGGAACAAGTTTTCTTGTCCGCCCGCCC |
| 42 | GCTCCCAAGATCCTCTTTTGAGGAACAAGTTTTCTTGTTCCAACTACG |
| 43 | AGCTTTTTAACTGCAGCAACTTTAATATACGCTATTGG |
| 44 | AGCTGGAATTACCGCGGCTGCTGGCACCAGACTTGCCC |
| 45 | TCCAATGGATCCTCGTTAAAGGATTTAAAGTGGACTCA |
| 46 | TTCCAATTACAGGGCCTCGAAAGAGTCCTGTATTGTTA |
| 47 | TTTTTCGTCACTACCTCCCCGGGTCGGGAGTGGGTAAT |
| 48 | TTGCGCGCCTGCTGCCTTCCTTGGATGTGGTAGCCGTT |
| 49 | TCTCAGGCTCCCTCTCCGGAATCGA |
| 50 | ACCCTGATTCCCCGTCACCCGTGGT |
| 51 | CACCATGGTATCCTCTTTTGAGGAACAAGTTTTCTTGTGGCACGGCGA |
| 52 | CTACCATCGATCCTCTTTTGAGGAACAAGTTTTCTTGTAAGTTGATAG |
| 53 | GGCAGACGTTTCCTCTTTTGAGGAACAAGTTTTCTTGTCGAATGGGTC |
| 54 | GTCGCCGCCATCCTCTTTTGAGGAACAAGTTTTCTTGTCGGGGGGCGT |
| 55 | GCGATCGGCCTCCTCTTTTGAGGAACAAGTTTTCTTGTCGAGGTTATC |
| 56 | TAGAGTCACCTCCTCTTTTGAGGAACAAGTTTTCTTGTAAAGCCGCCG |
| 57 | GCGCCCGCCCCCCGGCCGGGGCCGGAGAGGGGCTGACC |
| 58 | GGGTTGGTTTTGATCTGATAAATGCACGCATCCCCCCC |
| 59 | GCGAAGGGGGTCAGCGCCCGTCGGCATGTATTAGCTCT |
| 60 | AGAATTACCACAGTTATCCAAGTGGGAGAGGAGCGAGC |
| 61 | GACCAAAGGAACCATAACTGATTTAATGAGCCATTCGC |
| 62 | AGTTTCACTGTACCGGCCGTGCGTACTTAGACATGCAT |
| 63 | GGCTTAATCTTTGAGACAAGCATAT |
| 64 | GCTACTGGCAGGATCAACCAGGTA |

### Table S9

**Table S9.** Mice NR_003278.3 18S rRNA oligonucleotide sequences for ID `3232`.

| Oligonucleotide Number | Sequence (5′→3′) |
|---|---|
| 1 | TTAATGATCCTTCCGCAGGTTCACCTACGGAAACCTTG |
| 2 | TTACGACTTTTACTTCCTCTAGATAGTCAAGTTCGACC |
| 3 | GTCTTCTCAGCGCTCCGCCAGGGCCGTGGGCCGACCCC |
| 4 | GGCGGGGCCGATCCGAGGGCCTCACTAAACCATCCAAT |
| 5 | CGGTAGTAGCGACGGGCGGTGTGTACAAAGGGCAGGGA |
| 6 | CTTAATCAACGCAAGCTTATGACCCGCACTTACTGGGA |
| 7 | ATTCCTCGTTCATGGGGAATAATTGCAATCCCCGATCC |
| 8 | CCATCACGAATGGGGTTCAACGGGTTACCCGCGCCTGC |
| 9 | CGGCGCAGGGTAGGCACACGCTGAGCCAGTCAGTGTAG |
| 10 | CGCGCGTGCAGCCCCGGACATCTAAGGGCATCACAGAC |
| 11 | CTGTTATTGCTCAATCTCGGGTGGCTGAACGCCACTTG |
| 12 | TCCCTCTAAGAAGTTGGGGGACGCCGACCGCTCGGGGG |
| 13 | TCGCGTAACTAGTTAGCATGCCAGAGTCTCGTTCGTTA |
| 14 | TCGGAATTAACCAGACAAAT |
| 15 | CGCTCCACCAACTAAGAACGG |
| 16 | CCATGCACCACCACCCACGGAATCGtttggatatcactcattagtggt |
| 17 | AGAAAGAGCTATCAATCTGTCAATCtttggatatcactcattagtggt |
| 18 | CTGTCCGTGTCCGGGCCGGGTGAGGTTTCCCGTGTTGA |
| 19 | GTCAAATTAAGCCGCAGGCCCACTCCTGGTGGTGCCCT |
| 20 | TCCGTCAATTCCTTTAAGTTTCAGCTTTGCAACCATAC |
| 21 | TCCCCCCGGAACCCAAAGACTTTGGTTTCCCGGAAGCT |
| 22 | GCCCGGCGGGTCATGGGAATAAC |
| 23 | GCCGCCGCATCGCCAGTCGGCATCGtttggatatcactcattagtggt |
| 24 | TTTATGGTCGGAACTACGACGGTATtttggatatcactcattagtggt |
| 25 | CTGATCGTCTTCGAACCTCCGACTTtttggatatcactcattagtggt |
| 26 | TCGTTCTTGATTAATGAAAACATTCTTGGCAAATGCTT |
| 27 | TCGCTCTGGTCCGTCTTGCGCCGGTCCAAGAATTTCAC |
| 28 | CTCTAGCGGCGCAATACGAATGCCCCCGGCCGTCCCTC |
| 29 | TTAATCATGGCCTCAGTTCCGAAAACCAACAAAATAGA |
| 30 | ACCGCGGTCCTATTCCATTATTC |
| 31 | CTAGCTGCGGTATCCAGGCGGCTCGtttggatatcactcattagtggt |
| 32 | GGCCTGCTTTGAACACTCTAATTTTtttggatatcactcattagtggt |
| 33 | TTCAAAGTAAACGCTTCGGGCCCCGCGGGACACTCAGC |
| 34 | TAAGAGCATCGAGGGGGCGCCGAGAGGCAAGGGGCGGG |
| 35 | GACGGGCGGTGACTCGCCTCGCGGCGGACCGCCCGCCC |
| 36 | GCTCCCAAGATCCAACTACGAGCTTTTTAACTGCAGCA |

| | |
|---|---|
| 37 | ACTTTAATATACGCTATTGGAGC |
| 38 | TGGAATTACCGCGGCTGCTGGCACCtttggatatcactcattagtggt |
| 39 | AGACTTGCCCTCCAATGGATCCTCGtttggatatcactcattagtggt |
| 40 | TTAAAGGATTTAAAGTGGACTCATTtttggatatcactcattagtggt |
| 41 | CCAATTACAGGGCCTCGAAAGAGTCCTGTATTGTTATT |
| 42 | TTTCGTCACTACCTCCCCGGGTCGGGAGTGGGTAATTT |
| 43 | GCGCGCCTGCTGCCTTCCTTGGATGTGGTAGCCGTTTC |
| 44 | TCAGGCTCCCTCTCCGGAATCGAACCCTGATTCCCCGT |
| 45 | CACCCGTGGTCACCATGGTAGGCACGGCGACTACCATC |
| 46 | GAAAGTTGATAGGGCAGACGTTCGAATGGGTCGTCGCC |
| 47 | GCCACGGGGGGCGTGCGATCGGCCCGAGGTTATCTAGA |
| 48 | GTCACCAAGCCGCCGGCGCCCGACCCCCGGCCGGAGCC |
| 49 | GGGAGGGAGCTCACCGGGTTGGTTTTGATCTGATAAAT |
| 50 | GCACGCATCCCCCCCCGGGAAGGGGGGTCAGCGCCCGT |
| 51 | CGGCATGTATTAGCTCTAGAATTACCACAGTTATCCAA |
| 52 | GTAGGAGAGGAGCGAGCGACCAAAGGAACCATAACTGA |
| 53 | TTTAATGAGCCATTCGCAGTTTCACTGTACCGGCCGTG |
| 54 | CGTACTTAGACATGCATGGCTTAATCTTTGAGACAAGC |
| 55 | ATATGCTACCTGGCAGGATCAACCAGGT |

### Table S10

**Table S10.** *Salmonella* Typhimurium 16S rRNA oligonucleotide sequences for ID `3113`.

| Oligonucleotide Number | Sequence (5′→3′) |
|---|---|
| 1 | TAAGGAGGTGATCCAACCGCAGGTTNCCCTACGGTTAC |
| 2 | CTTGTTACGACTTCACCCCAGTCATGAATCACAAAGTG |
| 3 | GTAAGCGCCCTCCCGAAGGTTAAGCTACCTACTTCTTT |
| 4 | TGCAACCCACTCCCATGGTGTGACGGGCGGTGTGTACA |
| 5 | AGGCCCGGGAACGTATTCACCGTGGCATTCTGATCCAC |
| 6 | GATTACTAGCGATTCCGACTTCATGGAGTCGAGTTGCA |
| 7 | GACTCCAATCCGGACTACGACGCACTTTATGAGGTCCG |
| 8 | CTTGCTCTCGCGAGGTCGCTTCTCTTTGTATGCGCCAT |
| 9 | TGTAGCACGTGTGTAGCCCTGGTCGTAAGGGCCATGAT |
| 10 | GACTTGACGTCATCCCCACCTTCCTC |
| 11 | CAGTTTATCACTGGCAGTCTCCTTTGA |
| 12 | GTTCCCGACCTAATCGCTGGCAACAtttggatatcactcattagtggt |
| 13 | AAGGATAAGGGTTGCGCTCGTTGCGtttggatatcactcattagtggt |
| 14 | GGACTTAACCCAACATTTCACAACAtttggatatcactcattagtggt |
| 15 | CGAGCTGACGACAGCCATGCAGCACCTGTCTCACAGTT |
| 16 | CCCGAAGGCACCAATCCATCTCTGGATTCTTCTGTGGA |
| 17 | TGTCAAGACCAGGTAAGGTTCTTCGCGTTGCATCGAAT |
| 18 | TAAACCACATGCTCCACCGCTTGTGCGGCCCCCGTCAA |
| 19 | TTCATTTGAGTTTTAACCTTGCG |
| 20 | GCCGTACTCCCCAGGCGGTCTACTTtttggatatcactcattagtggt |
| 21 | AACGCGTTACGTCCGGAAGCCACGCCTCAAGGGCACAA |
| 22 | CCTCCAAGTAGACATCGTTTACGGCGTGGACTACCAGG |
| 23 | GTATCTAATCCTGTTTGCTCCCCACGCTTTCGCACCTG |
| 24 | AGCGTCAGTCTTTGTCCAGGGGGCCGCCTTCGCCACCG |
| 25 | GTATTCCTCCAGATCTCTACGCA |
| 26 | TTTCACCGCTACACCTGGAATTCTAtttggatatcactcattagtggt |
| 27 | CCCCCCTCTACAAGACTCAAGCCTGCCAGTTTCGAATG |
| 28 | CAGTTCCCAGGTTGAGCCCGGGGATTTCACATCCGACT |
| 29 | TGACAGACCGCCTGCGTGCGCTTTACGCCCAGTAATTC |
| 30 | CGATTAACGCTTGCACCCTCCGTATTACCGCGGCTGCT |
| 31 | GGCACGGAGTTAGCCGGTGCTTC |
| 32 | TTCTGCGGGTAACGTCAATTGCTGCtttggatatcactcattagtggt |
| 33 | GGTTATTAACCACAACACCTTCCTCtttggatatcactcattagtggt |
| 34 | CCCGCTGAAAGTACTTTACAACCCGtttggatatcactcattagtggt |
| 35 | AAGGCCTTCTTCATACACGCGGCATGGCTGCATCAGGC |
| 36 | TTGCGCCCATTGTGCAATATTCCCCACTGCTGCCTCCC |

| | |
|---|---|
| 37 | GTAGGAGTCTGGACCGTGTCTCAGTTCCAGTGTGGCTG |
| 38 | GTCATCCTCTCAGACCAGCTAGGGATCGTCGCCTTGGT |
| 39 | GAGCCGTTACCTCACCAACTAGCTAATCCCATCTGGGC |
| 40 | ACATCTGATGGCAAGAGGCCCGAAGGTCCCCCTCTTTG |
| 41 | GTCTTGCGACGTTATGCGGTATTAGCCACCGTTTCCAG |
| 42 | TAGTTATCCCCCTCCATCAGGCAGTTTCCCAGACATTA |
| 43 | CTCACCCGTCCGCCACTNNNNAGCGAAGNGCAAGCTGC |
| 44 | TTCCTGTTACCGTTCGACTTGCATGTGTTAGGCCTGCC |
| 45 | GCCAGCGTTCAATCTGAGC |
| 46 | CATGATCAAACTCTTCAATT |

**Table S11**

**Table S11.** MS2 rRNA oligonucleotide sequences for ID `111`.

| Oligonucleotide Number | Sequence (5′→3′) |
|---|---|
| 1 | CACTCCGTTCCCTACAACGAGCCTAAATTCATATGACT |
| 2 | CGTTATAGCGGACCGCGTGTCTGATCCACGGCGCACAT |
| 3 | TGGTCTCGGACCAATAGAGCCGCTCTCAGAGCGCGGGG |
| 4 | GGTAACGGTTGCTTGTTCAGCGAACTTCTTGTAAGGCG |
| 5 | CTGCATCCTGCAACTTGTGCCCCATAGGAGCACCGTTG |
| 6 | GAGAACGTGCATTGCCCAAACAACGACGATCGGTAGCC |
| 7 | AGAGAGGAGGTTGCCAATAAGGCTACGGATGCTGGTTT |
| 8 | GTAAAACATCCGGATCCCATGACAAGGATTTGTCATGT |
| 9 | AAGAAACCTTCTCTATTTATCTGACCGCGATCACCATT |
| 10 | CGCCTCCCGTTCCTCTTTTGAGGAACAAGTTTTCTTGTAGCTTAGCGA |
| 11 | TAGCTAAGGTTCCTCTTTTGAGGAACAAGTTTTCTTGTACGACGGGTC |
| 12 | GCCTCGTCATTCCTCTTTTGAGGAACAAGTTTTCTTGTTACCAGAACC |
| 13 | TAAGGTCGGATCCTCTTTTGAGGAACAAGTTTTCTTGTTGCTTTGTGA |
| 14 | GCAATTCGTCTCCTCTTTTGAGGAACAAGTTTTCTTGTCCTTAAGTAA |
| 15 | GCAATTGCTGTCCTCTTTTGAGGAACAAGTTTTCTTGTTAAAGTCGTC |
| 16 | ACTGTGCGGATCACCGCTTCCAGTAGCGACAG |
| 17 | AAGCAATTGATTGGTAAATTTCGAGAGAAAGATCGCGA |
| 18 | GGAAGATCAATACATAAAGAGTTGAACTTCTTTGTTGT |
| 19 | CTTCGACATGGGTAATCCTCATGTTTGAATGGCCGGCG |
| 20 | TCTATTAGTAGATGCCGGAGTTTGCTGCGATTGCTGAG |
| 21 | GGAATCGGGTTTCCATCTTTTAGGAGACCTTGCATTGC |
| 22 | CTTAACAATAAGCTCGCAGTCGGAATTCGTAGCGAAAA |
| 23 | TTGGAATGGTTAGTTCCATATTTAAGTACGAACGCCAT |
| 24 | GCGGCTACAGGAAGCTCTACACCACCAACAGTCTGGGT |
| 25 | TGCCACTTTAGGCACCTCGACTTTGATGGTGTATTTGC |
| 26 | GATTCTGCGCAGAGCTCTGACGAACGCTACAGGTTACT |
| 27 | TTGTAAGCCTGTGAACGCGAGTTAGAGCTGATCCATTC |
| 28 | AGCGACCCCGTTAGCGAAGTTGCTTGGGGCGACAGTCA |
| 29 | CGTCGCCAGTTCCTCTTTTGAGGAACAAGTTTTCTTGTTCCGCCATTG |
| 30 | TCGACGAGAATCCTCTTTTGAGGAACAAGTTTTCTTGTCGAACTGAGT |
| 31 | AAAGTTAGAATCCTCTTTTGAGGAACAAGTTTTCTTGTGCCATGCTTC |
| 32 | AAACTCCGGTTCCTCTTTTGAGGAACAAGTTTTCTTGTTGAGGGCTCT |
| 33 | ATCTAGAGAGTCCTCTTTTGAGGAACAAGTTTTCTTGTCCGTTGCCTG |
| 34 | ATTAATGCTATCCTCTTTTGAGGAACAAGTTTTCTTGTACGCATCTAA |
| 35 | GGTATGGACCATCGAGAAAGGAGACTTTACGT |
| 36 | ACGCGCCAGTTGTTGGCCATACGGATTGTACCCCTCGA |

| | |
|---|---|
| 37 | TGCATGGCTGAGATTTGGGCCTTAGCAGTGCCCTGTCT |
| 38 | CTCCACAGTCCACCCGTAGGGAGCGTCAACGCTTATGA |
| 39 | TGGACTCACCCGTTATTACGTCAGTAACTGTTCCTGAC |
| 40 | ATGTAGGAGCATCCCACGGGGGCCGTAAGGCCCTCGAG |
| 41 | CATGTTACCTACAGGTAGGAGCCAGTCGACAACGAATG |
| 42 | AGAAAGGCACCTTTTCCCACACTATACCTAGTGGGTTC |
| 43 | AAGATACCTAGAGACGACAACCATGCCAAACGTGCATC |
| 44 | GTTTATGTAAAACCATATCACGATACGTCGCGATATGT |
| 45 | TGCACGTTGTTCCTCTTTTGAGGAACAAGTTTTCTTGTCTGGAAGTTT |
| 46 | GCAGCTGGATTCCTCTTTTGAGGAACAAGTTTTCTTGTACGACAGACG |
| 47 | GCCATCTAACTCCTCTTTTGAGGAACAAGTTTTCTTGTTTGATGTTAG |
| 48 | TACCGACCTGTCCTCTTTTGAGGAACAAGTTTTCTTGTACGTACGGCT |
| 49 | CTCATAGGAATCCTCTTTTGAGGAACAAGTTTTCTTGTGAAACTCTTG |
| 50 | AAGGTGAACCTCCTCTTTTGAGGAACAAGTTTTCTTGTTTCGTAAGCA |
| 51 | TCTCATATGCACCCTGGATATCACTCATTAGT |
| 52 | GGTAACCAACCGAACTGCAACTCCAACCACCTGCCGGC |
| 53 | CACGTGTTTTGATCGAAACTTTCGATCTTCGTTTAGGG |
| 54 | CAAGGTAGCGGAGCGCCTGGCGCCAATTACCGCGACGA |
| 55 | GCGGCAGTGTACGCCTTCACGAGCGCAATGGTTTGCGT |
| 56 | CGCGAGTTGTGAGGCTGTCGACCTGGCCTCTGCTAAAG |
| 57 | CAACACCAAGGTTAAAATTACCCTGGGTGACCTTTTGC |
| 58 | AGGACTTCGGTCGACGCCCGGTTCGCAACGTTCTGCGG |
| 59 | CACTTCGATGTAAGTCAAGTTTTGGCTTACAGGGAAGA |
| 60 | GGCTGTAGCAGGAGCGTGCGTCGAGGGAGAAGCCGAAA |
| 61 | CCGGCTTTCTCCTCGTACGGGCGACCCCACGATGACCCACTTCGCTTGTAG |
| 62 | GCACCTTGATCTATCGATGTGACACTTAACGCCCCCCGTGAATACGGAGA |
| 63 | GGGGTAGTGCCACTGTTTCGTTTTGGCCCCAGTCGAGTTAAAACGACCGG |
| 64 | GAGTCCAGTTCGAACGATATTTTAAAGAGAATGAGTTATCTTCAGTCTCA |
| 65 | CCGTCCGCGTAAACGCGAACGGAGGGGACGAAGGTCTCGTTCTCCCTATC |
| 66 | AAGGGTACTAAAAGCTCGCACAGGTCAAACCTCCTAGGAATGGAATTCCG |
| 67 | GCTACCTACAGCGATAGCCATGGTAGCGTCTCGCTAAAGACATTAAAAAT |
| 68 | GGCATTAGCTCGACAGGAAGTTGAGCAGGACCCCGAAAGGGGTCCCACCC |
| 69 | TGGGTGGTAACTAGCCAAGCAGCTAGTTACCAAATCGGGAGAATCCCGGG |
| 70 | TCCTCTCTTTAGGGGGAGGTCCCTGGGCCGAAGCCCGCCCACCTTTCGGT |
| 71 | GGAGCCGGACCGCTTTCGCACCCGTGCTCTTTCGAGCACACCCACCCCGT |
| 72 | TTACGGGGGTCCCTCGGTCAGCTACCGAGGAGAGCTCGCTGGCCCACACT |
| 73 | CCTGAGGGAATGTGGGAACCGGCGTTAGCCACTCCGAAGTGCGTATAACG |
| 74 | CGCACGCCGGCGGACTTCATGCTGTCGGTGATTTCACCTCCAGTATGGAA |
| 75 | CCACGCTATGTAGCGACCACTGTCGTGCTTTTCGCTGAAGAACTTGCGTT |
| 76 | CTCGAGCGATACGAGCAAGACGGAAACCCGAGGTACGGGTATCCGCGAGC |
| 77 | AGCCGCCCGTACGGAGTCTTGGTGTATACCGAGACTGCCGTAGGCGGGCT |

| | |
|---|---|
| 78 | GACTACGTAGTAGTCGGCAGCGAGGTCCGTCCCACCGAAGAACATCGAAG |
| 79 | GCACCTGGGAGGAGAGCCGTACCCACACCTTATAGAGGCGTGGATCTGAC |
| 80 | ATACCTCCGACAACTCCCCAACCCCGTAGCCGATTTAATATCAGCATCAG |
| 81 | GGCGAAGAGATTGTCAACAGGTTTCTTGATGTAAAACGGTTTGACATCGA |
| 82 | CACCACGGTAAAAGTGCGCGCCGCAGCTCTCGCGAAAGAGCCCGGACACG |
| 83 | AACGTTTTACGAAGATTCGGTTTAAAACCGTAGTAGGCAAGTGCCTCTAG |
| 84 | CACACGGGGTGCAATCTCACTGGGACATATAATATCGTCCCCGTAGATGC |
| 85 | CTATGGTTCCGGCGTTACCAAAATGGATTTGGGTCGCTTTGACTATTGCC |
| 86 | CAGAATATCATGGACTCTAGCTCAAATGTGAACCCATTTCCCATTGTGGA |
| 87 | AAATAGTTCCCATCGTATCGTCTCGCCATCTACGATTCCGTAGTGTGAGC |
| 88 | GGATACGATCGAGATATGAATATAGCTCTGGTGGGAGAAAACTCCACACC |
| 89 | AGGCGATCGGAGATGGAATCGGATGCAGACGATAAGTCTATCGTCGCAAG |
| 90 | CGAACCATCTACGCTGCCCTGCTGAGCCAGACGCTGGTTGATCGATTGAT |
| 91 | CATTCAGGTCTATACCAACGGATTTGAGCCGGCGTCTGATGAAAGCACCG |
| 92 | ACCCCTTTCTGGAGGTACATATTCATATCAGGCTCCTTAC |
| 93 | AGGCAGCCCGATCTATTTTATTATTCTTCGGAACTGTAAA |

**Table S12**

**Table S12.** M13 DNA oligonucleotide sequences for ID `111111`.

| Replaced oligonucleotides from Table S1. | Sequence (5′→3′) |
|---|---|
| **For the first bit:** 26-30 | ACATCACTTGTCCTCTTTTGAGGAACAAGTTTTCTTGTCCTGAGTAGA |
| | AGAACTCAAATCCTCTTTTGAGGAACAAGTTTTCTTGTCTATCGGCCT |
| | TGCTGGTAATTCCTCTTTTGAGGAACAAGTTTTCTTGTATCCAGAACA |
| | ATATTACCGCTCCTCTTTTGAGGAACAAGTTTTCTTGTCAGCCATTGC |
| | AACAGGAAAATCCTCTTTTGAGGAACAAGTTTTCTTGTACGCTCATGG |
| | AAATACCTACTCCTCTTTTGAGGAACAAGTTTTCTTGTATTTTGACGC |
| | TCAATCGTCTTCCTCTTTTGAGGAACAAGTTTTCTTGTGAAATGGATT |
| | ATTTACATTGTCCTCTTTTGAGGAACAAGTTTTCTTGTGCAGATTCAC |
| | CAGTCACACGACCAGTAATAAAAGGGACAT |
| **For the second bit:** 52-57 | TTACCTGAGCAAAAGAAGATGATGAAACAAACATCAAGAAAACA |
| | AAATTAATTATCCTCTTTTGAGGAACAAGTTTTCTTGTCATTTAACAA |
| | TTTCATTTGATCCTCTTTTGAGGAACAAGTTTTCTTGTATTACCTTTT |
| | TTAATGGAAATCCTCTTTTGAGGAACAAGTTTTCTTGTCAGTACATAA |
| | ATCAATATATTCCTCTTTTGAGGAACAAGTTTTCTTGTGTGAGTGAAT |
| | AACCTTGCTTTCCTCTTTTGAGGAACAAGTTTTCTTGTCTGTAAATCG |
| | TCGCTATTAATCCTCTTTTGAGGAACAAGTTTTCTTGTTTAATTTTCC |
| | CTTAGAATCCTCCTCTTTTGAGGAACAAGTTTTCTTGTTTGAAAACAT |
| | AGCGATAGCTTCCTCTTTTGAGGAACAAGTTTTCTTGTTAGATTAAGA |
| | CGCTGAGAAGAGTCAATAGTGAAT |
| **For the third bit:** 79-85 | TGAATCTTACCAACGCTAACGAGCGTCTTTCCAGAGCCTAATTTGCCAGT |
| | TACAAAATAATCCTCTTTTGAGGAACAAGTTTTCTTGTACAGCCATAT |
| | TATTTATCCCTCCTCTTTTGAGGAACAAGTTTTCTTGTAATCCAAATA |
| | AGAAACGATTTCCTCTTTTGAGGAACAAGTTTTCTTGTTTTTGTTTAA |
| | CGTCAAAAAATTCCTCTTTTGAGGAACAAGTTTTCTTGTGAAAATAGCA |
| | GCCTTTACAGTCCTCTTTTGAGGAACAAGTTTTCTTGTAGAGAATAAC |
| | ATAAAAACAGTCCTCTTTTGAGGAACAAGTTTTCTTGTGGAAGCGCAT |

| | |
|---|---|
| | TAGACGGGAGTCCTCTTTTGAGGAACAAGTTTTCTTGTAATTAACTGA |
| | ACACCCTGAATCCTCTTTTGAGGAACAAGTTTTCTTGTCAAAGTCAGA |
| | GGGTAATTGAGCGCTAATATCAGAGAGATAACCCACAAGAATTGAGTTAAGCCCAA |
| **For the fourth bit:** | TCACAAACAAATAAATCCTCATTAAAGCCAGAATGGAAAGCGCAGTCTCTGAATTT |
| 106-112 | ACCGTTCCAGTCCTCTTTTGAGGAACAAGTTTTCTTGTTAAGCGTCAT |
| | ACATGGCTTTTCCTCTTTTGAGGAACAAGTTTTCTTGTTGATGATACA |
| | GGAGTGTACTTCCTCTTTTGAGGAACAAGTTTTCTTGTGGTAATAAGT |
| | TTTAACGGGGTCCTCTTTTGAGGAACAAGTTTTCTTGTTCAGTGCCTT |
| | GAGTAACAGTTCCTCTTTTGAGGAACAAGTTTTCTTGTGCCCGTATAA |
| | ACAGTTAATGTCCTCTTTTGAGGAACAAGTTTTCTTGTCCCCCTGCCT |
| | ATTTCGGAACTCCTCTTTTGAGGAACAAGTTTTCTTGTCTATTATTCT |
| | GAAACATGAATCCTCTTTTGAGGAACAAGTTTTCTTGTAGTATTAAGA |
| | GGCTGAGACTCCTCAAGAGAAGGATTAGGATTAGCGGGGTTTTGCTCAGT |
| **For the fifth bit:** | CAAAGTACAACGGAGATTTGTATC |
| 134-139 | ATCGCCTGATTCCTCTTTTGAGGAACAAGTTTTCTTGTAAATTGTGTC |
| | GAAATCCGCGTCCTCTTTTGAGGAACAAGTTTTCTTGTACCTGCTCCA |
| | TGTTACTTAGTCCTCTTTTGAGGAACAAGTTTTCTTGTCCGGAACGAG |
| | GCGCAGACGGTCCTCTTTTGAGGAACAAGTTTTCTTGTTCAATCATAA |
| | GGGAACCGAATCCTCTTTTGAGGAACAAGTTTTCTTGTCTGACCAACT |
| | TTGAAAGAGGTCCTCTTTTGAGGAACAAGTTTTCTTGTACAGATGAAC |
| | GGTGTACAGATCCTCTTTTGAGGAACAAGTTTTCTTGTCCAGGCGCAT |
| | AGGCTGGCTGTCCTCTTTTGAGGAACAAGTTTTCTTGTACCTTCATCA |
| | AGAGTAATCTTGACAAGAACCGGATATTCATTACCCAAATCAAC |
| **For the sixth bit:** | ACAGTTGATTCCCAATTCTGCGAACGAGTA |
| 161-165 | GATTTAGTTTTCCTCTTTTGAGGAACAAGTTTTCTTGTGACCATTAGA |
| | TACATTTCGCTCCTCTTTTGAGGAACAAGTTTTCTTGTAAATGGTCAA |
| | TAACCTGTTTTCCTCTTTTGAGGAACAAGTTTTCTTGTAGCTATATTT |
| | TCATTTGGGGTCCTCTTTTGAGGAACAAGTTTTCTTGTCGCGAGCTGA |
| | AAAGGTGGCATCCTCTTTTGAGGAACAAGTTTTCTTGTTCAATTCTAC |
| | TAATAGTAGTTCCTCTTTTGAGGAACAAGTTTTCTTGTAGCATTAACA |
| | TCCAATAAATTCCTCTTTTGAGGAACAAGTTTTCTTGTCATACAGGCA |

AGGCAAAGAATCCTCTTTTGAGGAACAAGTTTTCTTGTTTAGCAAAAT

### Table S13

**Table S13.** *Salmonella* Typhimurium 23S rRNA oligonucleotide sequences for ID `3113`.

| Oligonucleotide Number | Sequence (5′→3′) |
|---|---|
| 1 | TTAAGCCTCACGGTTCATTAGTACCGGTTAGCTCAACG |
| 2 | CATCGCTGCGCTTACACACCCGGCCTATCAACGTCGTC |
| 3 | GTCTTCAACGTTCCTTCAGGAGACCTAAAGTCTCAGGG |
| 4 | AGAACTCATCTCGGGGCAAGTTTCGTGCTTAGATGCTT |
| 5 | TCAGCACTTATCTCTTCCGCATTTAGCTACCGGGCAGT |
| 6 | GCCATTGGCATGACAACCCGAACACCAGTGATGCGTCC |
| 7 | ACTCCGGTCCTCTCGTACTAGGAGCAGCCCCCCTCAGT |
| 8 | TCTCCAGCGCCCACGGCAGATAGGGACCGAACTGTCTC |
| 9 | ACGACGTTCTAAACCCAGCTCGCGTACCACTTTAAATG |
| 10 | GCGAACAGCCATACCCTTGGGACCTACTTCAGCCCCAG |
| 11 | GATGTGATGAGCCGACATCGAGGTGCCAAACACCGCCG |
| 12 | TCGATATGAACTCTTGGGCGGTATCAGCCTGTTATCCC |
| 13 | CGGAGTACCTTTTATCCGTTGAGCGATGGCCCTTCCAT |
| 14 | TCAGAACCACCGGATCACTATGACCTGCTTTCGCACCT |
| 15 | GCTCGCGCCGTCACGCTCGCAGTCAAGCTGGCTTATGC |
| 16 | CATTGCACTAACCTCCTGATGTCCGACCAGGATTAGCC |
| 17 | AACCTTCGTGCTCCTCCGTTACTCTTTAGGAGGAGACC |
| 18 | GCCCCAGTCAAACTACCCACCAGACACTGTCCGCAACC |
| 19 | CGGATTACGGGTCCACGTTAGAACATCAAACATTAAAG |
| 20 | GGTGGTATTTCAAGGTCGGCTCCATGCAGACTGGCGTC |
| 21 | CACACTTCAAAGCCTCCCACCTATCCTACACATCAAGG |
| 22 | CTCAATGTTCAGTGTCAAGCTATAGTAAAGGTTCACGG |
| 23 | GGTCTTTCCGTCTTGCCGCGGGTACACTGCATCTTCAC |
| 24 | AGCGAGTTCAATTTCACTGAGTCTCGGGTGGAGACAGC |
| 25 | CTGGCCATCATTACGCCATTCGTGCAGGTCGGAACTTA |

| | |
|---|---|
| 26 | CCCGACAAGGAATTTCGCTACCTTAGGACCGTTATAGT |
| 27 | TACGGCCGCCGTTTACCGGGGCTTCGATCAGGAGCTTC |
| 28 | GCTTGCGCTGACCCCATCAATTAACCTTCCGGCACCGG |
| 29 | GCAGGCGTCACACCGTATACGTCCACTTTCGT |
| 30 | GTTTGCACAGTGCTGTGTTTTTAATtttggatatcactcattagtggt |
| 31 | AAACAGTTGCAGCCAGCTGGTATCTtttggatatcactcattagtggt |
| 32 | TCGACTGACTTCAGCTCCGTGAGTAtttggatatcactcattagtggt |
| 33 | AATCACTTCACCTACGTGTCAGCGTGCCTTCTCCCGAA |
| 34 | GTTACGGCACCATTTTGCCTAGTTCCTTCACCCGAGTT |
| 35 | CTCTCAAGCGCCTTGGTATTCTCTACCTGACCACCTGT |
| 36 | GTCGGTTTGGGGTACGATTTGATGTTACCTGATGCTTA |
| 37 | GAGGCTTTTCCTGGAAGCAGGGC |
| 38 | ATTTGTTGCTTCAGCACCGTAGTGCtttggatatcactcattagtggt |
| 39 | CTCGTCATCACGCCTCAGTGTTAAAGTGAACCGGATTT |
| 40 | ACCTGGAACACACACCTACACGCTTAAACCGGGACAAC |
| 41 | CGTCGCCCGGCCAACATAGCCTTCTCCGTCCCCCCTTC |
| 42 | GCAGTAACACCAAGTACGGGAATATTAACCCGTTTCCC |
| 43 | ATCGACTACGCCTTTCGGCCTCG |
| 44 | CCTTAGGGGTCGACTCACCCTGCCCtttggatatcactcattagtggt |
| 45 | CGATTAACGTTGGACAGGAACCCTTGGTCTTCCGGCGA |
| 46 | GCGGGCTTTTCACCCGCTTTATCGTTACTTATGTCAGC |
| 47 | ATTCGCACTTCTGATACCTCCAGCAACCCTCACAGGTC |
| 48 | ACCTTCGCAGGCTTACAGAACGCTCCCCTACCCAACAA |
| 49 | CGCATCAGCGTCGCTGCCGCAGC |
| 50 | TTCGGTGCATGGTTTAGCCCCGTTAtttggatatcactcattagtggt |
| 51 | CATCTTCCGCGCAGGCCGACTCGACtttggatatcactcattagtggt |
| 52 | CAGTGAGCTATTACGCTTTCTTTAAtttggatatcactcattagtggt |
| 53 | ATGATGGCTGCTTCTAAGCCAACATCCTGGCTGTCTGG |
| 54 | GCCTTCCCACATCGTTTCCCACTTAACCATGACTTTGG |

| | |
|---|---|
| 55 | GACCTTAGCTGGCGGTCTGGGTTGTTTCCCTCTTCACG |
| 56 | ACGGACGTTAGCACCCGCCGTGTGTCTCCCGTGATAAC |
| 57 | ATTCTCCGGTATTCGCAGTTTGCATCGGGTTGGTAAGC |
| 58 | CGGGATGGCCCCCTAGCCGAAACAGTGCTCTACCCCCG |
| 59 | GAGATGAATTCACGAGGCGCTACCTAAATAGCTTTCGG |
| 60 | GGAGAACCAGCTATCTCCCGGTTTGATTGGCCTTTCAC |
| 61 | CCCCAGCCACAGGTCATCCGCTAATTTTTCAACATTAG |
| 62 | TCGGTTCGGTCCTCCAGTTAGTGTTACCCAACCTTCAA |
| 63 | CCTGCCCATGGCTAGATCACCGGGTTTCGGGTCTATAC |
| 64 | CCTGCAACTTAACGCCCAATTAAGACTCGGTTTCCCTC |
| 65 | CGGCTCCCCTATTCGGTTAACCTTGCTACAGAATATAA |
| 66 | GTCGCTGACCCATTATACAAAAGGTACGCAGTCACCCC |
| 67 | ACCCCAAAGCATTCACCGCTTGTTTTGTGTGTTGAATG |
| 68 | CTTTGGTGGTGGGGCTCCCACTGCTTGTACGTACACGG |
| 69 | TTTCAGGTTCTTTTTCACTCCCCTCGCCGGGGTTCTTT |
| 70 | TCGCCTTTCCCTCACGGTACTGGTTCACTATCGGTCAG |
| 71 | TCAGGAGTATTTAGCCTTGGAGGATGGTCCCCCCATAT |
| 72 | TCAGACAGGATACCACGTGTCCCGCCCTACTCATCGAG |
| 73 | CTCACAGCACATGCGCTTTTGTGTACGGGGCTGTCACC |
| 74 | CTGTATCGCGCGCCTTTCCAGACGCTTCCACTAACACA |
| 75 | CATGCTGATTCAGGCTCTGGGCTCCTCCCCGTTCGCTC |
| 76 | GCCGCTACTGGGGGAATCTCGGTTGATTTCTTTTCCTC |
| 77 | GGGGTACTTAGATGTTTCAGTTCCCCCGGTTCGCCTCA |
| 78 | TTAACCTATGGATTCAGTTAATGATAGTGTGACGAGTC |
| 79 | ACACTGGGTTTCCCCATTCGGGTATCGCCGGTTATAAC |
| 80 | GGTTCATATCACCTTACCGGCGCTTATCGCAGATTAGC |
| 81 | ACGCCCTTCATCGCCTCTGACTGCCAGG |
| 82 | GCATCCACCGTGTACGCTTAGTCGCTTA |

### Table S14

**Table S14.** Complementary *A. baumannii* 16S rRNA oligonucleotide sequences for ID \`0331\`.

| **Oligonucleotide Number** | **Sequence (5′→3′)** |
|---|---|
| 1 | TAACTGAAGAGTTTGATCATGGCTCAGATTGAACGCTG |
| 2 | GCGGCAGGCTTAACACATGCAAGTCGAGCGGGGGAAGG |
| 3 | TAGCTTGCTACTGGACCTAGCGGCGGACGGGTGAGTAA |
| 4 | TGCTTAGGAATCTGCCTATTAGTGGGGGACAACATCTC |
| 5 | GAAAGGGATGCTAATACCGCATACGTCCTACGGGAGAA |
| 6 | AGCAGGGGATCTTCGGACCTTGCGCTAATAGATGAGCC |
| 7 | TAAGTCGGATTAGCTAGTTGGTGGGGTAAAGGCCTACC |
| 8 | AAGGCGACGATCTGTAGCGGGTCTGAGAGGATGATCCG |
| 9 | CCACACTGGGACTGAGACACGGCCCAGACTCCTACGGG |
| 10 | AGGCAGCAGTGGGGAATATTGGACAATGGGGGGAACCC |
| 11 | TGATCCAGCCATGCCGCGTGTGTGAAGAAGGCCTTATG |
| 12 | GTTGTAAAGCACTTTAAGCGAGGAGGAGGCTACTTTAG |
| 13 | TTAATACCTAGAGATAGTGGACGTTACTCGCAGAATAA |
| 14 | GCACCGGCTAACTCTGTGCCAGCAGCCGCGGTAATACA |
| 15 | GAGGGTGCGAGCGTTAATCGGATTTACTGGGCGTAAAG |
| 16 | CGTGCGTAGGCGGCTTATTAAGTC |
| 17 | GGATGTGAAATCCCCGAGCTTAACTtttggatatcactcattagtggt |
| 18 | TGGGAATTGCATTCGATACTGGTGAtttggatatcactcattagtggt |
| 19 | GCTAGAGTATGGGAGAGGATGGTAGtttggatatcactcattagtggt |
| 20 | AATTCCAGGTGTAGCGGTGAAATGCGTAGAGATCTGGA |
| 21 | GGAATACCGATGGCGAAGGCAGCCATCTGGCCTAATAC |
| 22 | TGACGCTGAGGTACGAAAGCATGGGGAGCAAACAGGAT |
| 23 | TAGATACCCTGGTAGTCCATGCCGTAAACGATGTCTAC |
| 24 | TAGCCGTTGGGGCCTTTGAGGCT |
| 25 | TTAGTGGCGCAGCTAACGCGATAAGtttggatatcactcattagtggt |

| | |
|---|---|
| 26 | TAGACCGCCTGGGGAGTACGGTCGCtttggatatcactcattagtggt |
| 27 | AAGACTAAAACTCAAATGAATTGACtttggatatcactcattagtggt |
| 28 | GGGGGCCCGCACAAGCGGTGGAGCATGTGGTTTAATTC |
| 29 | GATGCAACGCGAAGAACCTTACCTGGCCTTGACATACT |
| 30 | AGAAACTTTCCAGAGATGGATTGGTGCCTTCGGGAATC |
| 31 | TAGATACAGGTGCTGCATGGCTGTCGTCAGCTCGTGTC |
| 32 | GTGAGATGTTGGGTTAAGTCCCG |
| 33 | CAACGAGCGCAACCCTTTTCCTTACtttggatatcactcattagtggt |
| 34 | TTGCCAGCATTTCGGATGGGAACTTTAAGGATACTGCC |
| 35 | AGTGACAAACTGGAGGAAGGCGGGGACGACGTCAAGTC |
| 36 | ATCATGGCCCTTACGGCCAGGGCTACACACGTGCTACA |
| 37 | ATGGTCGGTACAAAGGGTTGCTACACAGCGATGTGATG |
| 38 | CTAATCTCAAAAAGCCGATCGTAGTCCGGATTGGAGTC |
| 39 | TGCAACTCGACTCCATGAAGTCGGAATCGCTAGTAATC |
| 40 | GCGGATCAGAATGCCGCGGTGAATACGTTCCCGGGCCT |
| 41 | TGTACACACCGCCCGTCACACCATGGGAGTTTGTTGCA |
| 42 | CCAGAAGTAGCTAGCCTAACTGCAAAGAGGGCGGTTAC |
| 43 | CACGGTGTGGCCGATGACTGGGGTGAAGTCGTAACAAG |
| 44 | GTAGCCGTAGGGGAACCTGCGGCTGGATCACCTCCTTA |

### Table S15

**Table S15.** Complementary *Salmonella* serovar 23S rRNA oligonucleotide sequences for IDs. The same set was used for all three serovar 13S rRNA ID assembly.

| Oligonucleotide Number | Sequence (5′→3′) |
|---|---|
| 1 | CTTCGGCGTTGTAAGGTTAAGCCTCACGGTTCATTAGT |
| 2 | ACCGGTTAGCTCAACGCATCGCTGCGCTTACACACCCG |
| 3 | GCCTATCAACGTCGTCGTCTTCAACGTTCCTTCAGGAG |
| 4 | ACTCTAAGTCTCAGGGAGAACTCATCTCGGGGCAAGTT |
| 5 | TCGTGCTTAGATGCTTTCAGCACTTATCTCTTCCGCAT |
| 6 | TTAGCTACCGGGCAGTGCCATTGGCATGACAACCCGAA |
| 7 | CACCAGTGATGCGTCCACTCCGGTCCTCTCGTACTAGG |
| 8 | AGCAGCCCCCCTCAGTTCTCCAGCGCCCACGGCAGATA |
| 9 | GGGACCGAACTGTCTCACGACGTTCTAAACCCAGCTCG |
| 10 | CGTACCACTTTAAATGGCGAACAGCCATACCCTTGGGA |
| 11 | CCTACTTCAGCCCCAGGATGTGATGAGCCGACATCGAG |
| 12 | GTGCCAAACACCGCCGTCGATATGAACTCTTGGGCGGT |
| 13 | ATCAGCCTGTTATCCCCGGAGTACCTTTTATCCGTTGA |
| 14 | GCGATGGCCCTTCCATTCAGAACCACCGGATCACTATG |
| 15 | ACCTGCTTTCGCACCTGCTCGCGCCGTCA |
| 16 | CGCTCGCAGTCAAGCTGGCTtttggatatcactcattagtggt |
| 17 | TATGCCATTGCACTAACCTCtttggatatcactcattagtggt |
| 18 | CTGATGTCCGACCAGGATTAtttggatatcactcattagtggt |
| 19 | GCCAACCTTCGTGCTCCTCCGTTACTCTTTAGGAGGAG |
| 20 | ACCGCCCCAGTCAAACTACCCACCAGACACTGTCCGCA |
| 21 | ACCCGGGTAACGGGTCCACGTTAGAACATCAAACATTA |
| 22 | AAGGGTGGTATTTCAAGGTCGGCTCCATGCAGACTGGC |
| 23 | GTCCACACTTCAAAGCCTCCCACCTATCCTACACATCA |
| 24 | AGGCTCAATGTTCAGTGTCAAGCTATAGTAAAGGTTCA |
| 25 | CGGGGTCTTTCCGTCTTGCCGCGGGTACACTGCATCTT |

| | |
|---|---|
| 26 | CACAGCGAGTTCAATTTCACTGAGTCTCGGGTGGAGAC |
| 27 | AGCCTGGCCATCATTACGCCATTCGTGCAGGTCGGAAC |
| 28 | TTACCCGACAAGGAATTTCGCTACCTTAGGACCGTTAT |
| 29 | AGTTACGGCCGCCGTTTACCGGGGCTTCGATCAGGAGC |
| 30 | TTCGCTTGCGCTGACCCCATCAATTAACCT |
| 31 | TCCGGCACCGGGCAGGCGTCtttggatatcactcattagtggt |
| 32 | ACACCGTATACGTCCACTTTCGTGTTTGCACAGTGCTG |
| 33 | TGTTTTTAATAAACAGTTGCAGCCAGCTGGTATCTTCG |
| 34 | ACTGACTTCAGCTCCATGAGTAAATCACTTCACCTACG |
| 35 | TGTCAGCGTGCCTTCTCCCGAAGTTACGGCACCATTTT |
| 36 | GCCTAGTTCCTTCACCCGAGTTCTCTCAAGCGCCTTGG |
| 37 | TATTCTCTACCTGACCACCTGTGTCGGTTTGGGGTACG |
| 38 | ATTTGATGTTACCTGATGCTTAGAGGCTTTTCCTGGAA |
| 39 | GCAGGGCATTTGTTGCTTCAGCAC |
| 40 | CGTAGTGCCTCGTCGTCACGCCTC |
| 41 | AGTGTTAAAGTGAACCGGATTTACCTGGAACACACACC |
| 42 | TACACGCTTAAACCGGGACAACCGTCGCCCGGCCAACA |
| 43 | TAGCCTTCTCCGTCCCCCCTTCGCAGTAACACCAAGTA |
| 44 | CGGGAATATTAACCCGTTTCCCATCGACTACGCCTTTC |
| 45 | GGCCTCGCCTTAGGGGTCGACTCACCCTGCCCCGATTA |
| 46 | ACGTTGGACAGGAACCCTTGGTCTTCCGGCGAGCGGGC |
| 47 | TTTTCACCCGCTTTATCGTTACTTATGTCAGCATTCGC |
| 48 | ACTTCTGATACCTCCAGCATGCCTCACGACACACC |
| 49 | TTCACAGGCTTACAGAACGCTCCCCTACCCAACAACAC |
| 50 | ACAGTGTCGCTGCCGCAGCTTCGGTGCATGGTTTAGCC |
| 51 | CCGTTACATCTTCCGCGCAGGCCGACTCGACCAGTGAG |
| 52 | CTATTACGCTTTCTTTAAATGATGGCTGCTTCTAAGCC |
| 53 | AACATCCTGGCTGTCTGGGCCTTCCCACATCGTTTCC |
| 54 | CACTTAACCATGACTTTGGGtttggatatcactcattagtggt |

| | |
|---|---|
| 55 | ACCTTAGCTGGCGGTCTGGGtttggatatcactcattagtggt |
| 56 | TTGTTTCCCTCTTCACGACGGACGTTAGCACCCGCCGT |
| 57 | GTGTCTCCCGTGATAACATTCTCCGGTATTCGCAGTTT |
| 58 | GCATCGGGTTGGTAAGCCGGGATGGCCCCCTAGCCGAA |
| 59 | ACAGTGCTCTACCCCCGGAGATGAATTCACGAGGCGCT |
| 60 | ACCTAAATAGCTTTCGGGGAGAACCAGCTATCTCCCGG |
| 61 | TTTGATTGGCCTTTCACCCCCAGCCACAGGTCATCCGC |
| 62 | TAATTTTTCAACATTAGTCGGT |
| 63 | TCGGTCCTCCAGTTAGTGTTtttggatatcactcattagtggt |
| 64 | ACCCAACCTTCAACCTGCCCtttggatatcactcattagtggt |
| 65 | ATGGCTAGATCACCGGGTTTCGGGT |
| 66 | CTATACCCTGCAACTTAACGCCCG |
| 67 | GTTAAGACTCGGTTTCCCTC |
| 68 | CGGCTCCCCTATTCGGTTAA |
| 69 | CCTTGCTACAGAATATAAGT |
| 70 | CGCTGACCCATTATACAAAA |
| 71 | GGTACGCAGTCACACCCAAA |
| 72 | GGGTGCTCCCACTGCTTGTACGTACACGGTTTCAGGTT |
| 73 | CTTTTTCACTCCCCTCGCCGGGGTTCTTTTCGCCTTTC |
| 74 | CCTCACGGTACTGGTTCACTATCGGTCAGTCAGGAGTA |
| 75 | TTTAGCCTTGGAGGATGGTCCCCCCATATTCAGACAGG |
| 76 | ATACCACGTGTCCCGCCCTACTCATCGAGCTCACAGCA |
| 77 | CATGCGCTTTTGTGTACGGGG |
| 78 | CTGTCACCCTGTATCGCGCGtttggatatcactcattagtggt |
| 79 | CCTTTCCAGACGCTTCCACTtttggatatcactcattagtggt |
| 80 | AACACACATGCTGATTCAGGtttggatatcactcattagtggt |
| 81 | CTCTGGGCTCCTCCCCGTTCGCTCGCCGCTACTGGGGG |
| 82 | AATCTCGGTTGATTTCTTTTCCTCGGGGTACTTAGATG |
| 83 | TTTCAGTTCCCCCGGTTCGCCTCATTAACCTATGGATT |

| | |
|---|---|
| 84 | CAGTTAATGATAGTGTGACGAATCACACTGGGTTTCCC |
| 85 | CATTCGGGTATCGCCGGTTATAACGGTTCATATCACCT |
| 86 | TACCGGCGCTTATCGCAGATTAGCACGCCCTTCATCGC |
| 87 | CTCTGACTGCCAGGGCATCCACCGTGT |
| 88 | ACGCTTAGTCGCTTAACCTCACAACCC |
